# Homeotic compartment curvature and tension control spatiotemporal folding dynamics

**DOI:** 10.1101/2022.11.19.517180

**Authors:** Aurélien Villedieu, Lale Alpar, Isabelle Gaugue, Amina Joudat, François Graner, Floris Bosveld, Yohanns Bellaïche

## Abstract

Shape is a conspicuous and fundamental property of biological systems entailing the function of organs and tissues. While much emphasis has been put on how tissue tension and mechanical properties drive shape changes, whether and how a given tissue geometry influences subsequent morphogenesis remains poorly characterized. Here, we explored how curvature, a key descriptor of tissue geometry, impinges on the dynamics of epithelial tissue invagination. We found that the morphogenesis of the fold separating the adult *Drosophila* head and thorax segments is driven by the invagination of the Deformed (Dfd) homeotic compartment. Dfd controls invagination by modulating actomyosin organization and in-plane epithelial tension via the Tollo and Dystroglycan receptors. By experimentally introducing curvature heterogeneity within the homeotic compartment, we established that a curved tissue geometry converts the Dfd-dependent in-plane tension into an inward force driving folding. Accordingly, the interplay between in-plane tension and tissue curvature quantitatively explains the spatiotemporal folding dynamics. Collectively, our work highlights how genetic patterning and tissue geometry provide a simple design principle driving folding morphogenesis during development.

## Introduction

During development, folding transforms epithelial cell sheets into complex three-dimensional (3D) structures. It underlies morphogenetic processes such as gastrulation, neurulation and organogenesis^1,2^. Furthermore, folding can morphologically delineate tissue compartments, and fold positions often correlate with selector or homeotic gene patterns as observed in vertebrate rhombomeres and in insect appendages or body segments^3–6^. As any morphogenetic process, folding could result from the complex interplay between cell and tissue mechanical stress, mechanical properties, and organ geometry. So far, numerous studies have explored how mechanical force production and tissue mechanical properties control the dynamics of tissue invaginations^7,8^. These studies have defined several mechanisms associated with fold formation including: (i) cell apical constriction for which the surface area or anisotropy of the constricting domain could modulate fold shape^9–18;^ (ii) apical-basal shortening of the cells^19–22;^ (iii) cell delamination^23,24^; and (iv) tissue buckling due to compressive stresses^11,25–31^. Despite these advances, whether and how epithelial folding dynamics is regulated by curvature, an intrinsic property of any epithelia, has remained unexplored during development. Here, by studying *Drosophila* adult cervix (neck) morphogenesis, we initially aimed to understand how the insect body plan is morphologically compartmentalized by deep invaginations. These tissue invaginations are much larger than the cell height and their positions may correlate with homeotic gene expression patterns^6^. We therefore envisioned that studying the inter-segmental invagination at the head-thorax region could provide insights into how large out-of-plane displacements are generated, and whether and how homeotic genes are tied up with morphogenesis. Here, we describe how, by taking advantage of our imaging setup, we serendipitously uncovered that body curvature combined with in-plane tension controlled by the *Deformed* (*Dfd*) homeotic gene promotes tissue invagination.

## Results

### The invagination of the pupal Dfd homeotic domain is associated with adult neck formation

As in all Diptera, the *Drosophila* adult is characterized by a head-thorax regionalization manifested by a deep inter-segmental invagination where the neck is located^32^ (Fig. 1a). In contrast, the *Drosophila* pupa does not harbor such a fold at 14 hours after pupa formation (14 hAPF, Fig. 1a). To understand when and how the neck fold is formed, we performed time-lapse imaging using the E-Cadherin::3xGFP (Ecad::3xGFP) adherens junction (AJ) marker, focusing on a region covering the dorsal part of the thorax and the head (Fig. 1b). Time-lapse imaging revealed that the presumptive neck tissue starts to gradually invaginate at 18 hAPF forming a fold that progressively deepens at the head-thorax interface. This invagination is accompanied by convergent tissue flows from the head and thorax towards the presumptive neck; the head and thorax tissues thus forming the flanks of the neck fold (Fig. 1b and Supplementary Video 1). Even prior to its folding, the presumptive neck region can be identified by the presence of cells elongated along the medial-lateral (ML) axis (Fig. 1b and Extended Data Fig. 1a). We found that these cells, hereafter referred to as neck cells, are marked by the expression of the homeotic gene *Dfd*, which also labels the adult neck (Fig. 1c,d, Extended Data Fig. 1b and Supplementary Video 2a). To quantitatively characterize the folding dynamics, we tracked neck folding on ML transverse sections along the neck fold using Ecad::3xGFP (Fig. 1e). This enabled us to follow the successive positions of the apical fold front along the ML axis, and to precisely quantify the stereotypical tissue deepening dynamics for 8 hours and up to 40µm below the initial tissue plane (Fig. 1f, Extended Data Fig. 1c-i and Supplementary Note). We conclude that neck folding is initiated during early pupal morphogenesis by the invagination of the Dfd homeotic compartment.

**Fig. 1.**
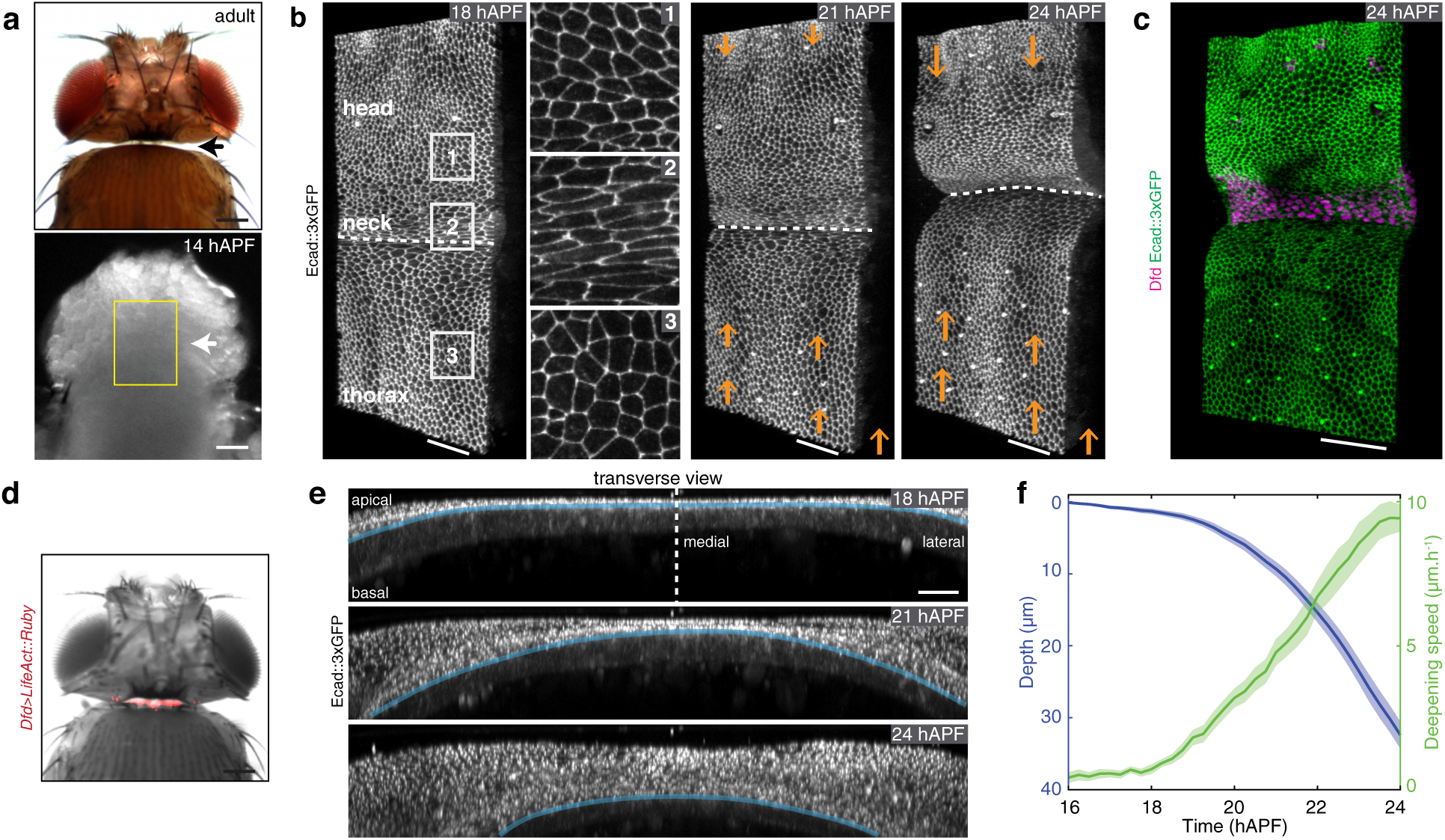
*Drosophila* neck folding during pupal development. **(a)** Dorsal view images of a *Drosophila* adult (top) and a 14 hAPF pupa (bottom). Arrows, position of the adult neck. Yellow box, region imaged in b. **(b)** Time-lapse 3D images of Ecad::3xGFP at 18, 21 and 24 hAPF during neck fold formation. See also Supplementary Video 1. 1-3 Close-ups of head, neck and thorax cells in the regions indicated in the left image. Orange arrows indicate the tissue velocity in the thorax and the head from 18 to 21 hAPF and from 21 to 24 hAPF (10µm.h^-1^, orange arrows in the bottom right). Dashed white line, apical fold front. **(c)** 3D image of Ecad::3xGFP and Dfd localizations in the dorsal neck region at 24 hAPF. **(d)** Dorsal view of a *Drosophila* adult showing *Dfd>LifeAct::Ruby* expression in the adult neck. Note some weak signal is also detected in very small regions of the head. **(e)** Transverse view time-lapse images of the neck region labelled by Ecad::3xGPF at 18, 21 and 24 hAPF. Blue line, position of the apical fold front. Dashed line, midline position. See also Supplementary Video 1 bottom. **(f)** Graph of the apical neck depth (blue, mean ± sem) and of neck deepening speed (green, mean ± sem) as a function of developmental time (*N*=10 pupae). Apical neck depth is defined relative to the initial position of the AJ labeled by Ecad::3xGFP at 16 hAPF. Scale bars, 50µm (a-d), 20µm (e).

### Apical and basal actomyosin dependent regulation of in-plane ML tension controls neck folding dynamics

To investigate whether and how the homeotic Dfd compartment invagination drives neck folding, we first characterized the organization of the actomyosin cytoskeleton and the mechanical stresses in the presumptive neck region. This revealed the presence of apical and basal actomyosin structures in the neck region (Supplementary Video 2b,c) and we further described these structures at 18 hAPF and 21 hAPF. In the apical domain, we observed the presence of supracellular apical Myosin light chain (MyoII) enrichment in the Dfd neck cells, as compared to the flanking tissues, as well as an apical actomyosin cable at the interface between the Dfd homeotic domain and the thorax cells marked by *Antennapedia* (*Antp*) expression (Fig. 2a and Supplementary Video 2). While we imaged MyoII and F-actin along the apical-basal axis of the neck cells, we noticed that MyoII and F-actin fibers are present basally, being well aligned with the ML axis from 21 hAPF onwards (Fig. 2b and Supplementary Video 2c). This basal network is present in the Dfd neck cells and in a few cells in the most anterior region of the thorax (Fig. 2b). Having identified these distinct apical and basal supracellular actomyosin structures, we then estimated in-plane tissue stresses along the anterior-posterior (AP) and ML axes by performing multi-photon laser ablation^33^ of these structures and measuring their recoil velocities (Fig. 2c,d, Extended Data Fig. 2a-f and Supplementary Video 3). This revealed that the apical and basal actomyosin structures are under tensile stress along the ML axis, while the stress is negligible along the AP axis (Extended Data Fig. 2a). The flanking head and thorax tissues also exhibited very little AP tension, and notably much lower ML tension compared to the neck region (Extended Data Fig. 2a). Furthermore, the neck ML apical and basal recoil velocities both increased during folding and were linearly correlated (Fig. 2c,d and Extended Data Fig. 2c). In addition, the apical and basal ML recoil velocities measured in the medial versus lateral neck regions were similar (Extended Data Fig. 2d-f).

**Fig. 2.**
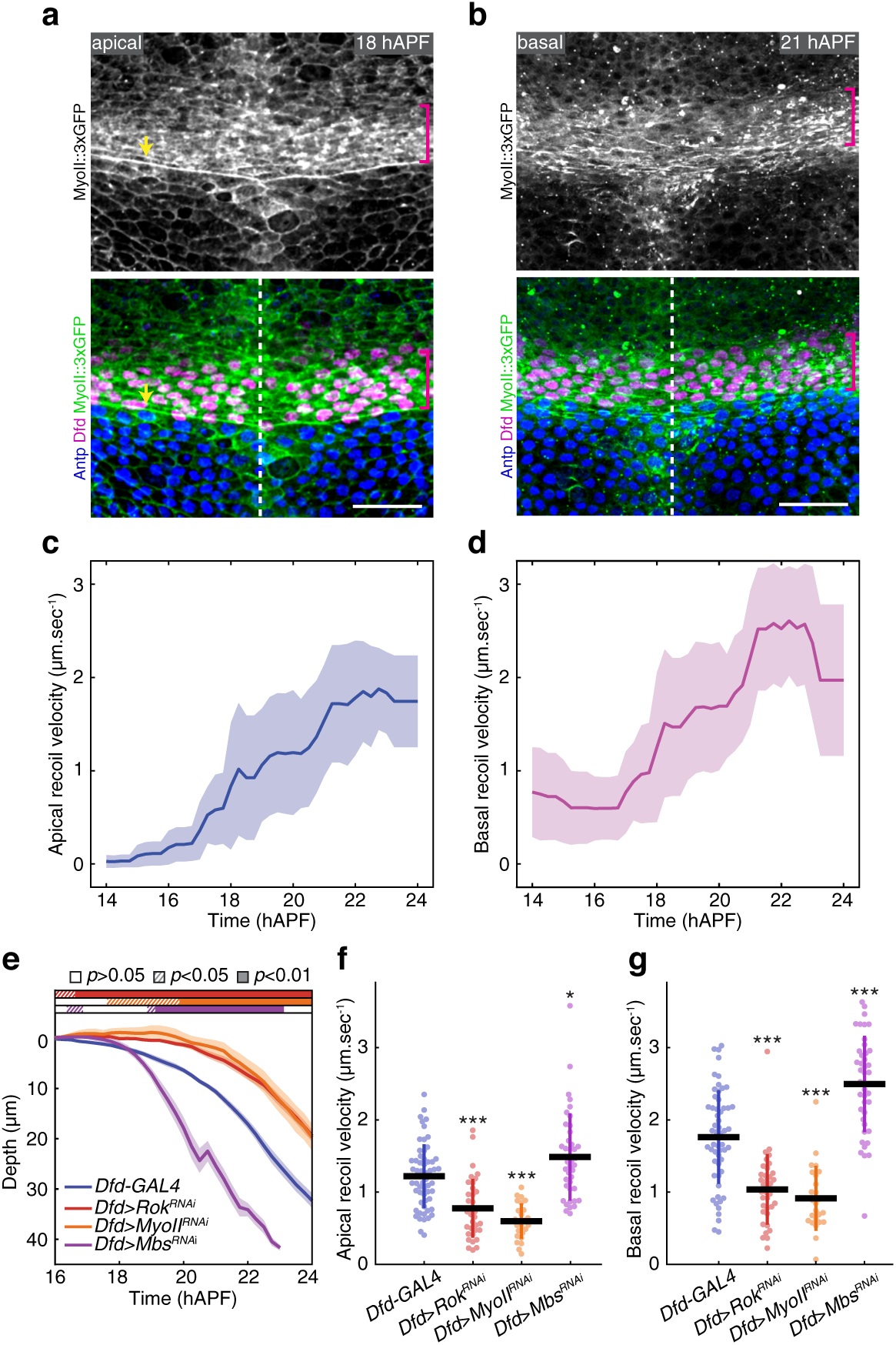
Dfd accumulation and MyoII-dependent tension during neck folding. **(a)** Apical MyoII::3xGFP localization (grey top, green bottom) as well as Dfd (magenta bottom) and Antp (blue bottom) nuclear distributions in the neck region at 18 hAPF. Magenta brackets, neck region. Yellow arrow, apical actomyosin cable at neck-thorax interface. Dashed line, midline position. **(b)** Basal MyoII::3xGFP localization (grey top, green bottom) as well as Dfd (magenta bottom) and Antp (blue bottom) nuclear distributions in the neck region at 21 hAPF. Magenta brackets, basal MyoII fibers oriented along the ML axis in the neck region. Dashed line, midline position. **(c,d)** Graph of the ML apical (c) and basal (d) initial recoil velocities (mean ± sd, averaged with a 2h sliding window) upon ablation in the medial neck region as a function of developmental time (*N=*127 pupae). See also Supplementary Video 3. **(e)** Graph of the apical neck depth (mean ± sem) in control *Dfd-GAL4* (*N=*9 pupae), *Dfd>Rok^RNAi^* (*N=*15 pupae), *Dfd>MyoII^RNAi^* (*N=*7 pupae) and *Dfd*>*Mbs^RNAi^* (*N=*13 pupae) tissues as a function of developmental time. Horizontal boxes: *p*-values of Welch tests performed between experimental condition and the *Dfd-GAL4* control at successive time points and color coded according to the experimental condition (white *p*>0.05, striped *p*<0.05, solid *p*<0.01). **(f)** Graph of the ML apical initial recoil velocities (mean ± sd) upon ablation in the neck region in control *Dfd-GAL4* (*N=*58 pupae), *Dfd*>*Rok^RNAi^* (*N=*36 pupae), *Dfd>MyoII^RNAi^* (*N=*25 pupae) and *Dfd>Mbs^RNAi^* (*N=*40 pupae). Ablations were performed at 20 ± 2 hAPF. *t*-test, * *p*<0.05, *** *p*<0.001. **(g)** Graph of the ML basal initial recoil velocities (mean ± sd) upon ablation in the neck region in control *Dfd-GAL4* (*N=*58 pupae), *Dfd>Rok^RNAi^* (*N=*36 pupae), *Dfd>MyoII^RNAi^* (*N=*25 pupae) and *Dfd>Mbs^RNAi^* (*N=*42 pupae). Ablations were performed at 20 ± 2 hAPF. *t*-test, *** *p*<0.001. Scale bars, 20µm.

Compromising MyoII activity in neck cells by either Rho kinase dsRNA (*Dfd>Rok^RNAi^*) or MyoII dsRNA (*Dfd>MyoII^RNAi^*) led to slower invagination as well as reduced apical and basal recoil velocities upon ablation (Fig. 2e-g and Extended Data Fig. 2g). In addition, injection of the Rok inhibitor (Y-27623) during folding completely abolished invagination (Extended Data Fig. 2h). Conversely, increasing MyoII activity by reducing MyoII phosphatase activity (*Dfd>Mbs^RNAi^*) caused a faster invagination and increased the apical and basal recoil velocities (Fig. 2e-g and Extended Data Fig. 2g). Taken together, our results indicate that neck folding is associated with the formation of apical and basal supracellular actomyosin structures and that MyoII activity within the Dfd domain promotes in-plane ML tension and folding dynamics.

### Dfd regulates in-plane tension via the Tollo and Dystroglycan receptors

We next aimed to identify the genetic regulators of the apical and basal actomyosin structures and mechanical stress within the Dfd homeotic compartment. Strongly reducing Dfd function in the neck (*Dfd>Dfd^RNAi^*) lowered fold deepening while diminishing apical and basal tensions (Fig. 3a-c and Extended Data Fig. 3a,c). In addition, reduced Dfd function lowered apical MyoII levels and the alignment of the basal actomyosin network in the neck (Fig. 3d and Extended Data Fig. 3d,e). In a survey of protein localization using functional GFP-tagged gene products, we uncovered that the Toll-like receptor Tollo was enriched at the level of the AJ in the neck cells whereas the extracellular matrix (ECM) receptor Dystroglycan (Dg) and its cytoplasmic adaptor Dystrophin (Dys) accumulated along the basal F-actin network (Fig. 3e,f, Extended Data Fig. 4a-c and Supplementary Video 2d,e). Tollo and Dys/Dg control tissue elongation by regulating actomyosin organization in the *Drosophila* embryo germband and oocyte, respectively^34–36^. However, their role in tissue folding is unexplored. We found that inhibiting Tollo function in the neck (*Dfd*>*Tollo^RNAi^*) slowed down neck deepening; a result that was further confirmed by analyzing *Tollo* null mutant animals (Fig. 3a and Extended Data Fig. 4d,e). Consistent with the notion that tension and deepening speed were reduced upon abrogating MyoII function, loss of Tollo function reduced apical tension and apical MyoII levels in the neck cells (Fig. 3b,g). In parallel with the analysis of Tollo function, we found that loss of Dg or Dys function (using *Dg* and *Dys* null mutant animals) as well as inhibiting Dg function in the neck (*Dfd*>*Dg^RNAi^*) also slowed down folding (Fig. 3h and Extended Data Fig. 4d,f). Accordingly, the loss of Dys or Dg function compromised the basal actomyosin organization in the neck and the most anterior cells of the thorax and led to a reduced basal tension (Fig. 3j-l and Extended Data Fig. 3d,e). We also analyzed neck folding in the double *Dys*,*Dfd>Tollo^RNAi^* mutant context, the effect of which on invagination dynamics was similar to the one observed in *Dfd>Dfd^RNAi^* as well as *Dfd>Tollo^RNAi^* or *Dys* mutant tissues (Extended Data Fig. 5a). This prompted us to test whether Tollo and Dys/Dg might also affect basal and apical tensions, respectively. Indeed, basal recoil velocity was reduced in *Dfd>Tollo^RNAi^* tissue, and conversely Dys and Dg loss of function affected apical tension (Fig. 3c,i). Since Tollo is mainly localized at the apical side of the cells, independently of Dg activity, and Dg is enriched at the basal side of the cells even in the absence of Tollo activity (Extended Data Fig. 5b,c), this could suggest that apical and basal tensions feedback on each other, or that Dg and Tollo have unforeseen indirect roles on the apical and basal tension, respectively. Lastly, we found that Dfd controls both Tollo and Dg distributions in the neck compartment (Fig. 3m,n). Altogether, we propose that Dfd regulates apical and basal actomyosin organizations and in-plane ML tension via Tollo and Dys/Dg to promote tissue folding.

**Fig. 3.**
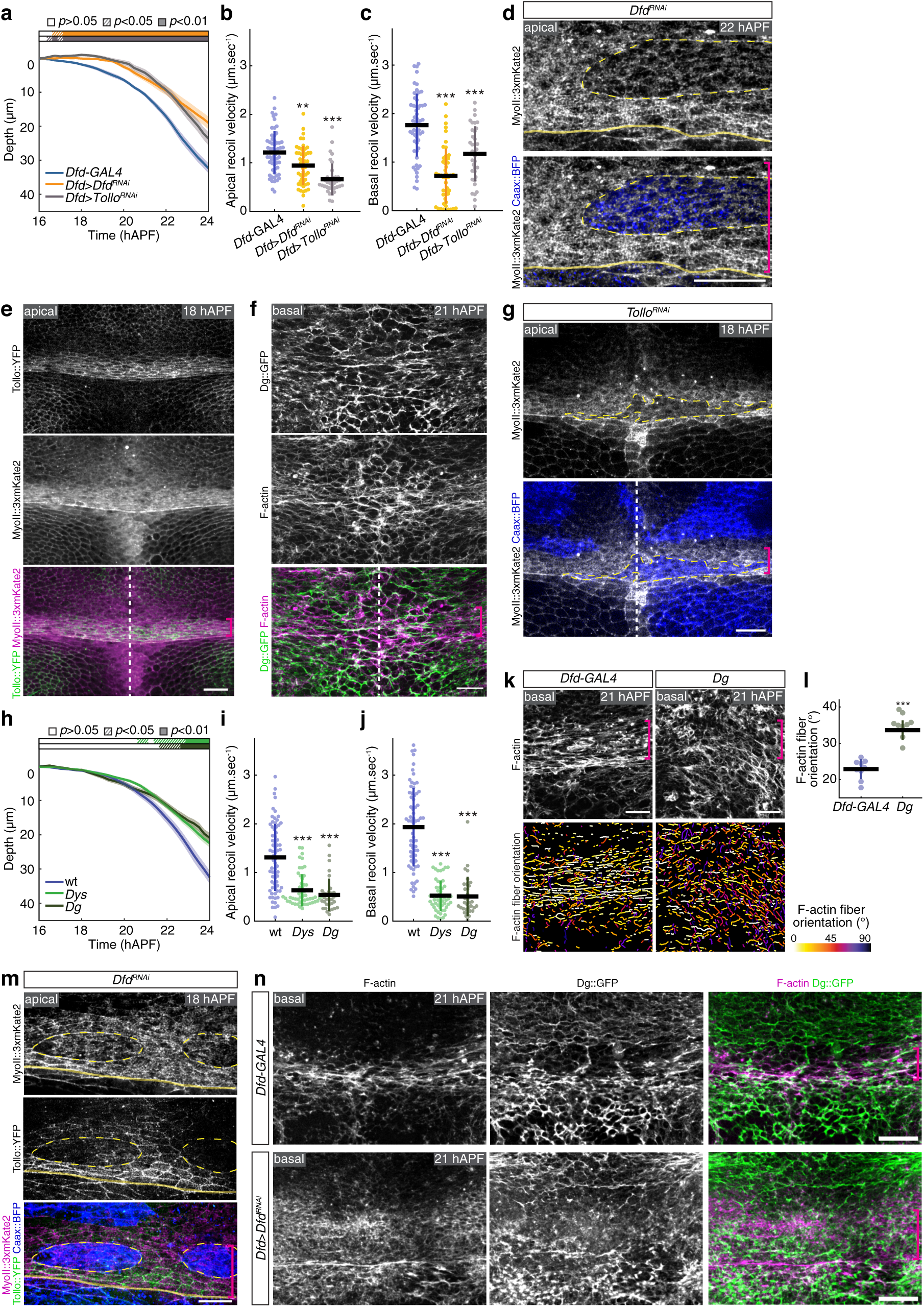
Control of ML in-plane tissue tension by Dfd, Tollo and Dys/Dg during neck folding. **(a)** Graph of the apical neck depth (mean ± sem) in control *Dfd-GAL4* (*N=*9 pupae), *Dfd>Dfd^RNAi^* (*N=*8 pupae) and *Dfd>Tollo^RNAi^* (*N=*11 pupae) tissues as a function of developmental time. Horizontal boxes: *p*-values of Welch tests performed between experimental condition and the *Dfd-GAl4* control at successive time points and color coded according to the experimental condition (white *p*>0.05, striped *p*<0.05, solid *p*<0.01). **(b,c)** Graph of the ML apical (b) and basal (c) recoil velocities after ablation (mean ± sd) in the neck region in *Dfd-GAL4* control (*N=*58 pupae), *Dfd>Dfd^RNAi^* (*N=*49 pupae) and *Dfd>Tollo^RNAi^* (*N=*36 pupae). Ablations were performed at 20 ± 2 hAPF. *t*-test, ** *p*<0.005, *** *p*<0.001. **(d)** Apical MyoII::3xmKate2 distribution in *Dfd^RNAi^* clones marked by Caax::BFP at 22 hAPF in the neck region. Yellow dashed lines, outlines a *Dg^RNAi^* clone in the neck region. Magenta bracket, neck region. Yellow line, neck-thorax boundary. **(e)** Apical Tollo::YFP (grey top, green bottom) and MyoII::3xmKate2 (grey center, magenta bottom) distributions at 18 hAPF in the neck region. Magenta bracket, neck region. Dashed line, midline position. **(f)** Basal Dg::GFP (grey top, green bottom) and F-actin (grey center, magenta bottom) distributions at 21 hAPF in the neck region. Magenta bracket, neck region. Dashed line, midline position. **(g)** Apical MyoII::3xmKate2 distribution in *Tollo^RNAi^* clones marked by Caax::BFP at 18 hAPF in the neck region. Yellow dashed lines, outlines a *Tollo^RNAi^* clone in the neck region. Magenta bracket, neck region. White dashed line, midline position. **(h)** Graph of the apical neck depth (mean ± sem) in wt (*N=*10 pupae), *Dys* (*N=*12 pupae) and *Dg* (*N=*10 pupae) tissues as a function of developmental time. Horizontal boxes: *p*-values of Welch tests performed between experimental condition and the wt control at successive time points and color coded according to the experimental condition (white *p*>0.05, striped *p*<0.05, solid *p*<0.01). **(i,j)** Graph of the ML apical (i) and basal (j) initial recoil velocities (mean ± sd) upon ablation in the neck region in wt (*N=*69 pupae), *Dys* (*N=*53 pupae) and *Dg* (*N=*33 pupae) tissues. Ablations were performed at 20 ± 2 hAPF. *t*-test, *** *p*<0.001. **(k)** Basal F-actin distribution (top) and segmentation of the F-actin fibers color-coded by orientation (bottom) at 21 hAPF in control *Dfd-GAL4* and *Dg* tissues. Segmented F-actin fibers are colored according to their orientation, ranging from 0° (along ML axis) to 90° (along AP axis). While F-Actin fiber orientation is affected in *Dg* mutant tissues, the phalloidin signal level is similar in control and *Dg* mutant tissues. Magenta brackets, neck region. **(l)** Graph of the F-actin fibers orientation relative to the ML axis (mean ± sd) in control *Dfd-GAL4* (*N=*9 pupae) and *Dg* (*N=*9 pupae) tissues in the neck region at 21 hAPF. Each F-actin fiber orientation is measured between 0° and 90° relative to the ML axis (i.e., 0° corresponding to a fiber parallel to the ML axis). Each dot corresponds to the fiber orientation value averaged for one animal. Welch test, *** *p*<0.001. **(m)** Apical MyoII::3xmKate2 (grey top, magenta bottom) and Tollo::YFP (grey center, green bottom) distributions in *Dfd^RNAi^* clones marked by Caax::BFP accumulation (blue bottom) at 18 hAPF. Yellow dashed lines, outlines of *Dfd^RNAi^* clones in the neck region. Magenta bracket, neck region. Yellow line, neck-thorax boundary. **(n)** Basal F-actin (grey left, magenta right) and Dg::GFP (grey center, green right) distributions in control *Dfd*>*GAL4* (top) and *Dfd*>*Dfd^RNAi^* (bottom) tissues at 21 hAPF. Magenta brackets, neck region. Scale bars: 10µm (d), 20µm (e-g, k,m,n).

### An interplay between in-plane tension and curvature can account for an inward force driving folding

Our results thus far put forward the homeotic control of actomyosin organization and the resulting in-plane tension contributing to neck folding. In addition, our analysis of mechanisms previously known^7,8^ to drive folding indicated that apical constriction associated with basal relaxation, apical-basal shortening, cell delamination or tissue buckling do not substantially contribute to neck folding (Extended Data Fig. 6 and Supplementary Note). We therefore sought for an alternative mechanism by which the in-plane ML tension in the neck region could generate a force moving cells towards the basal side of the epithelium, i.e. inside the animal. A hint was provided by the use of the coverslip needed to achieve high-resolution time-lapse imaging (Extended Data Fig. 7a,b). Without coverslip the neck region has a homogeneous curvature as observed on ML transverse section (Extended Data Fig. 7a). In contrast, the coverslip in contact with the apical ECM of the tissue locally flattened the tissue along the ML axis: the lateral regions being more curved than the medial one in contact with the coverslip (Fig. 1e and Extended Data Fig. 7b). Strikingly, while imaging with coverslip flattening, we noticed qualitatively that the lateral regions tended to invaginate faster than the flattened medial one, as was more evident at the beginning of the invagination (Supplementary Video 4). Interestingly, these disparities of invagination dynamics were not associated with differences in apical MyoII accumulation or recoil velocities between the medial and lateral neck regions (Extended Data Fig. 7c,d and Extended Data 2d,e).

To quantitatively investigate and model the role of tissue curvature, we first used time-lapse movies to determine both the curvature and deepening speed along the ML apical fold front (Fig. 4a). Confirming our initial qualitative observations, we found that at each given developmental time during tissue invagination, the deepening speed correlates well with curvature (Fig. 4b). We therefore aimed to describe neck folding dynamics based on simple physical considerations regarding a curved line under tension. If the line is straight, the tension does not generate any net resulting force perpendicular to the line. If the line is curved, the symmetry is broken and the line generates a net resulting force perpendicular to it, oriented towards the curvature center (Fig. 4c). The magnitude of the resulting force, per unit length of line, is the tension *T* multiplied by the curvature κ also called “Laplace” force; it has been applied previously to the one-dimensional border of a two-dimensional epithelium when modeling epithelium closure^37–41^. Here, since the tissue thickness (∼10µm) is smaller than the neck radius of curvature (∼300µm), we applied the equation of Laplace force to a thin epithelial strip which moves inwards perpendicular to its surface (see Supplementary Note). Inertia being negligible with respect to viscous energy dissipation, we write that the sum of this Laplace force and of the normal dissipative force is zero; the latter force being equal to the deepening speed (velocity *v_n_* of the line along the vector normal to the curve) multiplied by an unknown negative dissipative prefactor, −µ. So, the prediction is that *κT* − µ*v_n_*= 0 or equivalently, that *v_n_* is proportional to *κT* and is directed inwards. To experimentally test this prediction, we computed the product of the tissue ML tension and the apical fold front local curvature, and then compared it to the local deepening speed. Strikingly, we found that the data of folding dynamics at different developmental times now collapse along a single straight line showing a linear relationship with high correlation between the deepening speed on one hand, and the product of the apical fold front curvature and tissue ML tension on the other hand (Fig. 4d, R^2^=0.96). Interestingly, this proportionality relationship implies that the local modulation of neck folding dynamics with coverslip flattening is unlikely to be explained by changes in the friction or the adhesion between the apical ECM and the epithelium in the regions flattened by the coverslip (see Supplementary Note). Other mechanical changes indirectly induced by the flattening are equally unlikely due to the small magnitude of the resulting compression (see Supplementary Note). Importantly, we also assessed whether deepening speed is well predicted in experimental conditions where ML tension is modulated. We observed that in *Dfd>Rok^RNAi^*, *Dfd>MyoII^RNAi^, Dfd>Dfd^RNAi^*, *Dfd>Tollo^RNAi^*, *Dys*, or *Dg* mutant conditions for which the tensions were decreased, the deepening speed again linearly correlated with the product of the estimated tension and the local curvature (Fig. 4e,f, Extended Data Fig. 8 and Supplementary Note); this also holds true when increasing MyoII activity in *Dfd>Mbs^RNAi^* animals. Again, the data of folding dynamics collapse along a single straight line (Fig. 4e,f, Extended Data Fig. 8). We note that the slope of the linear relationship between the deepening speed and the product of estimated tissue ML tension and curvature varies between wild-type (wt) and some mutant conditions (Extended Data Fig. 8c). Since we assessed tissue tension via recoil velocity that depends upon tension as well as tissue mechanical properties (friction and viscosity)^33,42^, the slope variabilities may indicate that some mutant conditions could also affect tissue mechanical properties. This is for example the case for Dg loss of function exhibiting a higher slope than the one obtained in wt conditions (Extended Data Fig. 8c), as it promotes a strong decrease of recoil velocity while having a more modest impact on invagination dynamics (Fig. 2h-j). Last, while basal curvature measurements were less accurate, we also found a linear relationship between the deepening speed and the product of basal fold front curvature and ML tension on the basal fold front (Extended Data Fig. 9 and 10). Altogether, these results agree with a model in which the product of curvature and tension can account for neck folding dynamics.

**Fig. 4.**
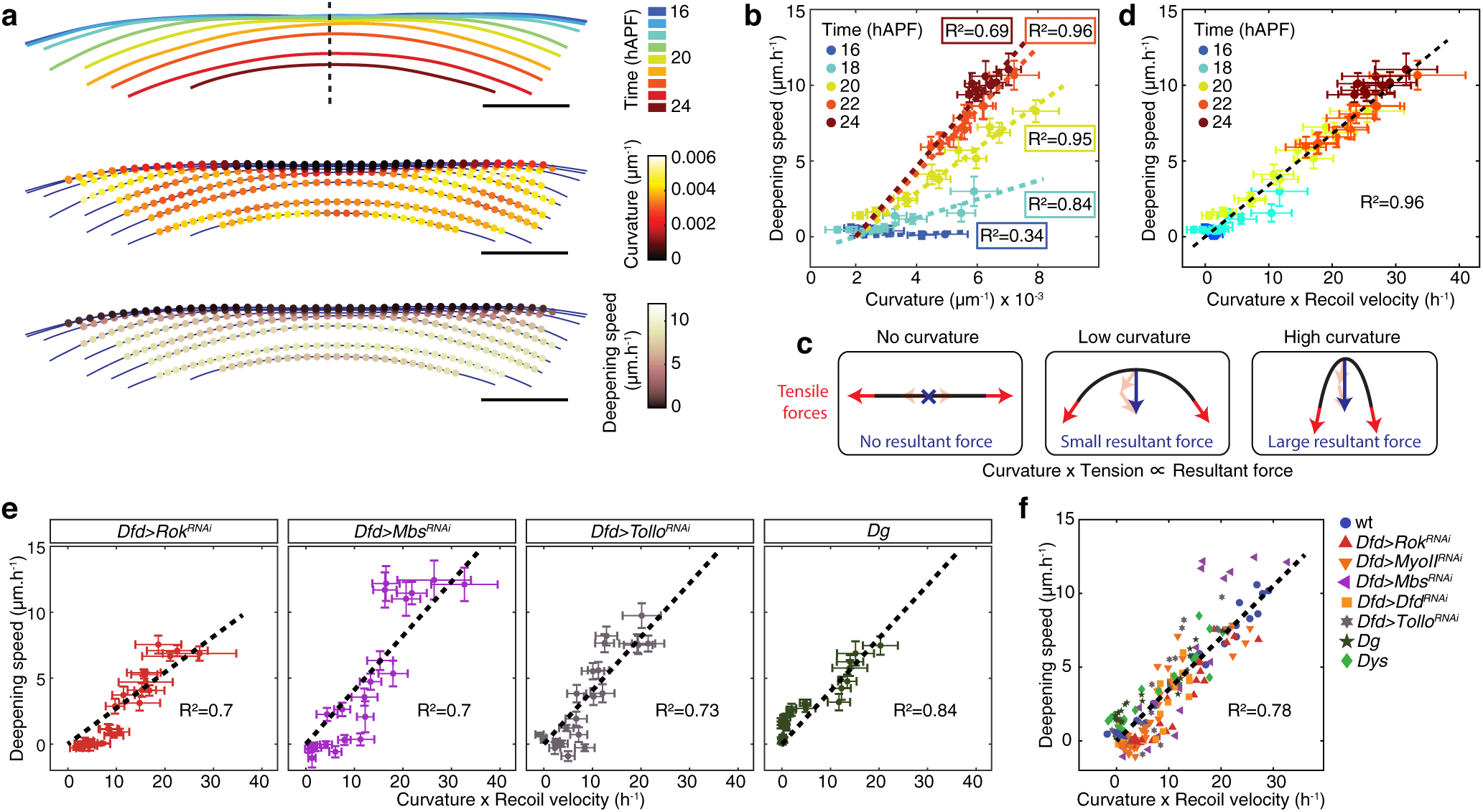
The product of local tissue curvature and ML tension can account for neck folding dynamics. **(a)** Successive positions of the apical fold front, color-coded according to time (top), local tissue curvature (middle) and local deepening speed (velocity along the vector normal to the curve) of the apical fold front (bottom) for an individual pupa. Dashed line, midline position. See also Supplementary Video 4. **(b)** Graph of the apical fold front deepening speed (mean ± sem) versus curvature (mean ± sem). Each point is the mean value (± sem) among 10 control animals for a given PositionML at a given color-coded developmental time. For each developmental time, a line passing through the origin was fitted (R^2^ values for each developmental time are indicated). **(c)** Schematic illustrating that the tissue line tension and the local tissue curvature define the magnitude of the normal resulting force (see text and Supplementary Note for details). **(d)** Graph of the apical fold front deepening speed (mean ± sem) and of the product of curvature and ML apical initial recoil velocity upon laser ablation in the neck region (mean ± sem) for a given PositionML at a color-coded developmental time. Datasets from Fig. 1f and Fig. 2c were used. A line passing through the origin was fitted to the average values (R^2^=0.96). **(e,f)** Graph of the apical fold front deepening speed (mean ± sem) versus the product of curvature and ML apical initial recoil velocity upon laser ablation in the neck region (mean ± sem) for a given PositionML and selected genotypes: *Dfd>Rok^RNAi^*, *Dfd>Mbs^RNAi^*, *Dfd>Tollo^RNAi^* and *Dg* (e), or all tested experimental conditions (f). Datasets from Fig. 2e, Fig. 3a, Fig. 3g and Extended Data Fig. 8 were used. A line passing through the origin was fitted (black dashed line, R^2^ values are indicated).

### Tissue curvature regulates the spatiotemporal dynamics of folding

Having investigated the role of in-plane tension, we next challenged the role of curvature by testing whether modulating the local tissue curvature would result in changes in deepening speed. Towards this goal, we developed a method to image the animal with a coverslip, but without inducing flattening, while maintaining a reasonable signal-to-noise ratio to track Ecad::3xGFP AJ signal (Fig. 5a and Supplementary Video 4). In conditions without flattening, the neck apical fold front shows a homogeneous and high curvature prior to folding. As with flattened controls, apical and basal tensions increase over time in non-flattened animals (Extended Data 11a,b). In agreement with a role of curvature in defining the resulting inward forces, we found that (i) tissue deepening speed is homogeneous along the ML axis in these animals, correlating with homogenously high curvature; (ii) despite the lower apical recoil velocities relative to the ones measured in flattened conditions, tissue deepening occurs earlier and faster in non-flattened animals (Fig. 5b-d and Extended Data 11a and Supplementary Video 4). Furthermore, the deepening speed is linearly correlated with the product of tension and curvature, suggesting that coverslip flattening modulates the invagination dynamics *per se*, rather than the underlying mechanisms (Fig. 5e and Extended Data Fig. 11c,d). If curvature modulates folding dynamics, reducing curvature in any region of the fold should locally reduce deepening speed. We thus flattened the lateral side of the neck region by mounting the pupa on its side; and used the contralateral non-flattened side as an internal control (Fig. 5f and Supplementary Video 4). In these experimental conditions, the flattened lateral side behaves like animals imaged with coverslip flattening in the medial domain, while the control non-flattened lateral side deepens with faster dynamics (Fig. 5g-i and Extended Data Fig. 11c). Altogether, we conclude that modulating the curvature led to changes in deepening speed that are predicted by the product of the curvature and the tension (Fig. 5e,j). Interestingly, we also noticed that independently of the initial curvature heterogeneity imposed by the coverslip, the tissue shape, with flat and curved regions, converged to a more homogenous curved geometry during invagination (Fig. 4a, Fig. 5g and Supplementary Video 4). In fact, since the velocity is proportional to curvature, a more curved region tends to shrink more quickly (Fig. 4a), thus with time the curvature increases and becomes more homogeneous (Extended Data Fig. 11e and Supplementary Note). This purely geometrical property may therefore account for a robust neck invagination process, independent of the initial heterogeneities in curvature due to our imaging set-up.

**Fig. 5.**
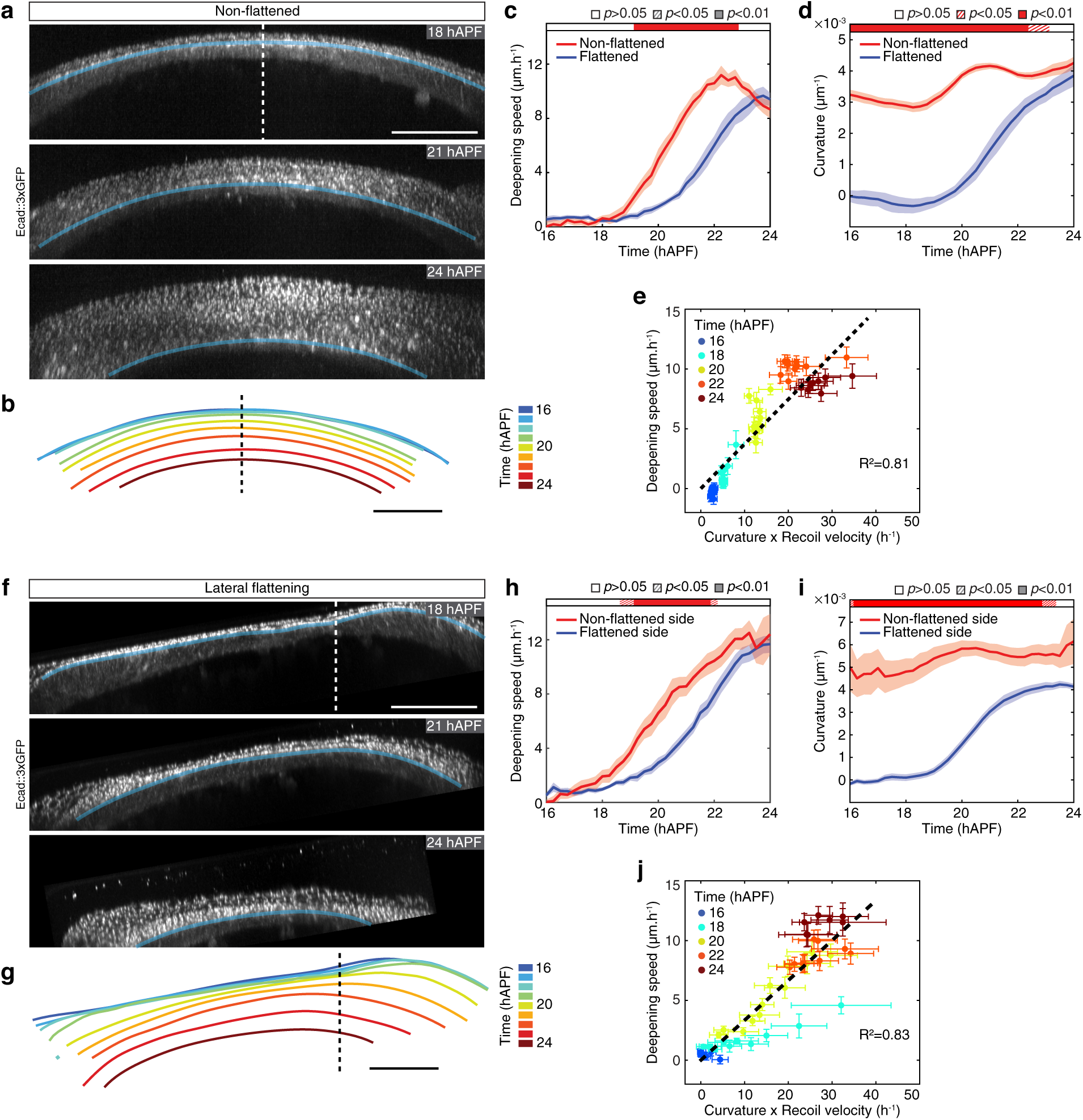
Local curvature along the neck fold controls folding dynamics. **(a)** Transverse view time-lapse images of the neck region labelled by Ecad::3xGPF at 18, 21 and 24 hAPF in a non-flattened pupa (i.e. imaged without coverslip flattening). Blue line, position of the apical fold front. Dashed line, midline position. See also Supplementary Video 4. **(b)** Successive positions of the apical fold front, color-coded according to time, in a pupa imaged without coverslip flattening (a). Dashed line, midline position. **(c)** Graph of apical fold front deepening speed (mean ± sem) in pupae imaged without coverslip flattening (non-flattened, *N=*11 pupae) and with coverslip flattening (flattened, *N=*101 pupae) as a function of developmental time. Horizontal box: *p-values* of Welch tests performed between non-flattened and flattened experimental condition at successive time points (white *p*>0.05, striped *p*<0.05, solid *p*<0.01). **(d)** Graph of curvature (mean ± sem) in pupae imaged without coverslip flattening (non-flattened, *N=*110 pupae) and with coverslip flattening (flattened, *N=*10 pupae) as a function of developmental time. Horizontal box: *p-values* of Welch tests performed between non-flattened and flattened experimental condition at successive time points (white *p*>0.05, striped *p*<0.05, solid *p*<0.01). **(e)** Graph of the apical fold front deepening speed (mean ± sem) and of the product of curvature and ML apical initial recoil velocity upon laser ablation in the neck region (mean ± sem) in pupae imaged without coverslip flattening for a given PositionML at a given color-coded developmental time. Datasets from Fig. 5c,d and Extended Data Fig. 11a were used. A line passing through the origin was fitted (R^2^=0.81). **(f)** Transverse view time-lapse images of the neck region labelled by Ecad::3xGPF at 18, 21 and 24 hAPF in a pupa flattened laterally relative to the midline. Blue line, position of the apical fold front. Dashed line, midline position. **(g)** Successive positions of the apical fold front, color-coded according to time, in a pupa laterally flattened (f). Dashed line, midline position. See also Supplementary Video 4. **(h)** Graph of apical fold front deepening speed (mean ± sem) of the non-flattened side and flattened side in pupae laterally flattened relative to the midline (as in (f), *N=*11) as a function of developmental time. Horizontal box: *p-values* of Welch tests performed between the non-flattened side and flattened side of the same pupae at successive time points (white *p*>0.05, striped *p*<0.05, solid *p*<0.01). **(i)** Graph of curvature (mean ± sem) of the non-flattened side and flattened side in pupae laterally flattened relative to the midline (as in (f), *N=*11) as a function of developmental time. Horizontal box: *p-values* of Welch tests performed between the non-flattened side and flattened side of the same pupae at successive time points (white *p*>0.05, striped *p*<0.05, solid *p*<0.01). **(j)** Graph of the apical fold front deepening speed (mean ± sem) and of the product of curvature and ML apical initial recoil velocity upon laser ablation in the neck region (mean ± sem) in laterally flattened pupae for a given PositionML at a given color-coded developmental time. Recoil velocities measured in flattened animals (Fig. 2c) were used for regions of initial curvatures below a threshold of 0.00015µm^-1^, while recoil velocities measured in non-flattened animals (Extended Data Fig. 11a) were used for regions of initial curvatures larger than 0.00015µm^-1^. A line passing through the origin was fitted (R^2^=0.83). Using a curvature threshold equal to 0 or varying it from 0.0005µm^-1^ to 0.002µm^-1^ leads to R^2^ varying from 0.81 to 0.83. Scale bars, 50µm.

Having found that Dfd homeotic compartment curvature impacts the spatiotemporal dynamics of neck invagination, we aimed at analyzing whether the presence of a curved region is necessary to promote folding. Prior to neck folding, we used a UV laser to ablate the tissue across its whole thickness to mechanically isolate the medial region from the lateral curved regions (Extended Data Fig. 11f). We then imaged neck folding while repeating the tissue severing ablations every 3h (Fig. 6a-g and Supplementary Video 5). When the isolated medial region was initially flattened by the coverslip, the tissue deepening speed was drastically reduced, while the tissue remained under ML tension (Fig. 6a-c and Extended Data Fig. 11g,h). Strikingly, without flattening, the isolated curved tissue initially deepened, and as it became flatter its deepening speed reduced (Fig. 6d-f). These experiments agree with the hypothesis that an initial curvature within the fold plane promotes folding and that a flat region can deepen if mechanically coupled to a curved one; thus, confirming the role of curvature in controlling the spatiotemporal dynamics of neck folding.

**Fig. 6.**
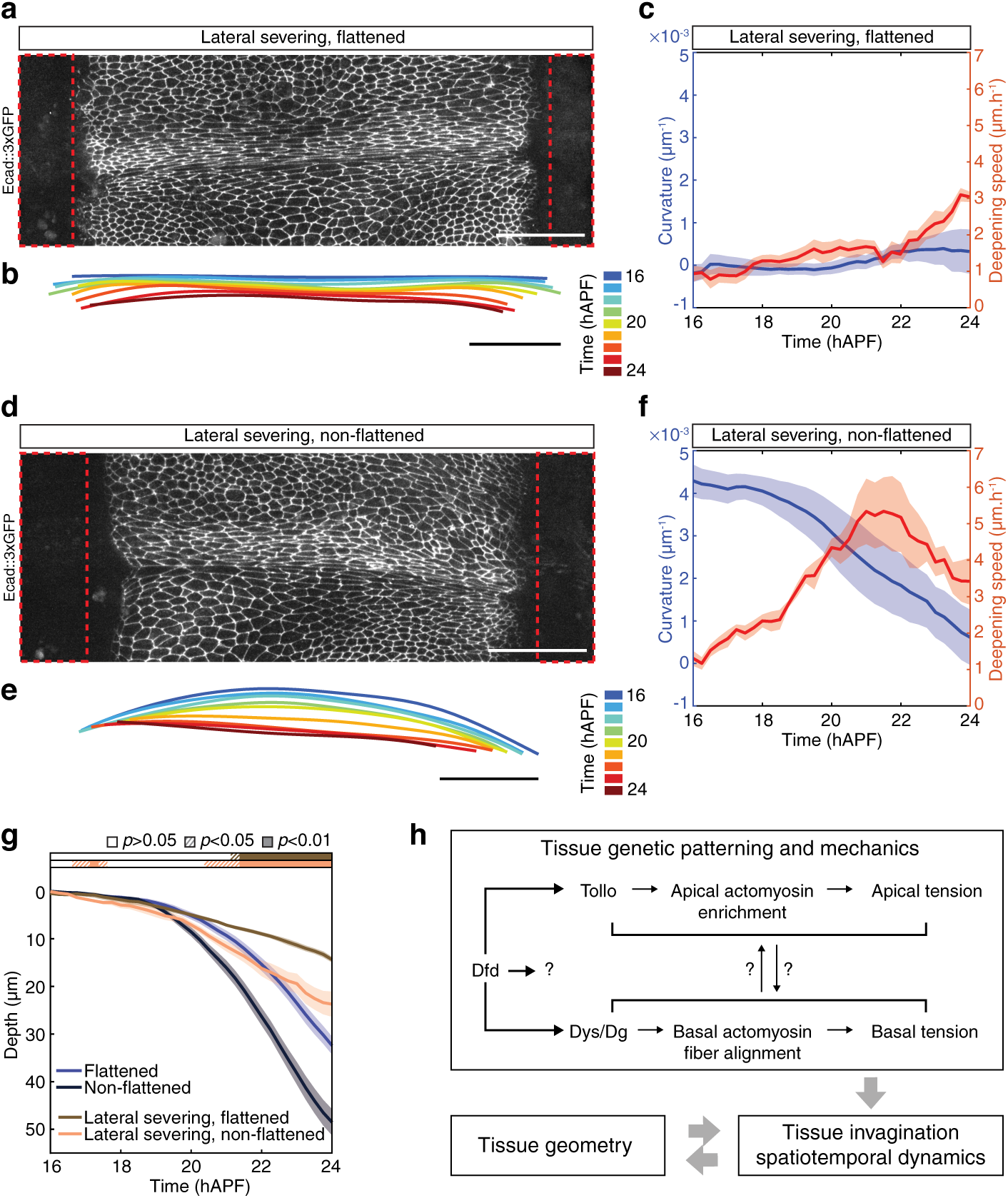
Local curvature is necessary for neck invagination. **(a,b)** Dorsal view image of Ecad::3xGFP distribution upon tissue severing ablation of each lateral side of the neck (red dashed boxes) (a) and corresponding successive positions (color-coded according to time) of the apical fold front in transverse views (b) in a pupa mounted with medial coverslip flattening. See also Supplementary Video 5. **(c)** Graph of curvature (blue, mean ± sem) and apical fold front deepening speed (red, mean ± sem) in pupae with two lateral side tissue severing ablations and imaged with medial coverslip flattening (*N=*9 pupae) as a function of developmental time. **(d,e)** Dorsal view image of Ecad::3xGFP distribution upon tissue severing ablation of each lateral side of the neck (red dashed boxes) (d) and corresponding color-coded successive positions of the apical fold front in transverse views (e) in a pupa mounted without coverslip flattening (non-flattened). See also Supplementary Video 5. **(f)** Graph of curvature (blue, mean ± sem) and apical fold front deepening speed (red, mean ± sem) in pupae with lateral side tissue severing ablations and imaged without coverslip flattening (*N=*6 pupae) as a function of developmental time. **(g)** Graph of apical neck depth (mean ± sem) in pupae without lateral side ablation and imaged with (flattened, *N=*10 pupae) and without (non-flattened, *N=*11 pupae) coverslip flattening as well as in pupae with two lateral side ablations and imaged with (flattened, *N=*9 pupae) and without coverslip flattening (non-flattened, *N*=6 pupae) as a function of developmental time. Horizontal boxes: *p*-values of Welch tests performed between the flattened and ablated, flattened condition (brown) as well as between the non-flattened and ablated, non-flattened condition (orange) at successive time points (white *p*>0.05, striped *p*<0.05, solid *p*<0.01). **(h)** Proposed model for the interplay between genetic patterning, tissue geometry and mechanical tension in the control of tissue folding. Genetic patterning leads to the establishment of in-plane ML apical and basal tension by Dfd-dependent regulation of the apical actomyosin enrichment and basal F-actin fiber organization via Tollo and Dys/Dg. Dfd might regulate Tollo and Dg accumulation via either similar or distinct mechanisms and our data do not exclude that additional Dfd downstream effectors modulate ML tension to promote neck invagination. The existence of a putative feedback between apical and basal tension is indicated by question marks (see discussion). Scale bars, 50µm.

## Discussion

The morphogenetic and functional specification of tissue compartments is a major trait of animal development. As such, the exploration of tissue morphogenesis and homeotic gene function is key to unravel the conserved mechanisms driving development. Here we uncovered that the function of Dfd homeotic gene in folding morphogenesis can be understood by integrating its activity with the initial animal geometry upon which it acts on. Accordingly, we propose that the establishment of in-plane tension combined with an initially convex curved geometry promotes an inward force controlling folding dynamics (Fig. 6h). Importantly, such spatiotemporal regulation by the product of tension and tissue curvature distinguishes this folding mechanism from known ones and defines a central role of tissue curvature in morphogenesis. While future work will be necessary to delineate how Dfd controls Tollo and Dg distributions and whether additional regulators act downstream of Dfd (Fig. 6h), the interplay between tissue tension and curvature likely expands beyond the fold we have explored here. Indeed, epithelial tissues *in vivo* are closed surfaces that by default harbor non-zero convex curvature and are often under tensile stress^7,43,44^. In particular, many tubular organs and curved embryonic epithelia are physically compartmentalized by transverse folds^3–6,22^, calling for further analyses of the roles of tissue and animal curvature during such fold formation. By linking in-plane tension and tissue curvature our work more generally helps understanding the role of tissue geometry in biology.

In addition to uncovering the critical role of tissue geometry in folding dynamics, our findings raise several important questions in the field of tissue morphogenesis. A distinctive feature of the folding process we have characterized is that the apical and basal tissue tensions are highly correlated. We found that apical Tollo and basal Dg regulate basal and apical tensions without controlling each other’s localizations and we did not detect any short time scale (tens of seconds) mechanical coupling between apical and basal tensions (Extended Data Fig. 6i,j). Therefore, we envision that the correlated apical and basal tensions are more likely linked to a long-time scale mechanical feedback or to an indirect role of Tollo or Dg on the basal or apical tension, respectively (Fig. 6h), via for example the apical-basal microtubule networks. The characterization of the putative mechanical feedback and the potential indirect roles of Tollo and Dg will be important to better understand how tissue folding and tissue thickness are controlled during neck morphogenesis, and more generally how the crosstalk between apical and basal tension is regulated in epithelial tissues. We note that we have used laser ablation to probe mechanical tension as validated in multiple experimental contexts^33,42^. Therefore, it remains difficult to fully explain small differences in apical and basal recoil velocities as they could reflect differences in the apical versus basal tissue mechanical properties, due to distinct friction on the apical versus basal ECM, for example. An additional novel feature unveiled by our work relates to the presence of two types of supracellular actomyosin structures in the apical region: the apical actomyosin cable at the boundary between neck and thorax and the strong actomyosin enrichment in the neck cells. We did not aim at separating the contributions of these two supracellular structures, and instead targeted to decipher the role of tissue curvature. Indeed, our theoretical model used to explore the role of tissue geometry gives very good predictions when treating the neck as a line and measuring recoil velocities across the two structures. We foresee that the characterization of the putative respective roles of these two actomyosin structures could help to delineate how (i) the boundary between homeotic compartments is controlled, (ii) the detailed sagittal shape of the invagination is modulated, and (iii) the feedback between apical and basal tensions is controlled. Finally, a last feature relates to the large scale of the neck fold we studied, and thus, the permissive versus active contributions of the neighboring tissue flows. Since the neck fold is very large relative to the size of the cells, the head and thorax tissue flows are substantial relative to previous folding processes driven by apical constriction^7,8^. Our ablation of head and thorax tissues illustrated that on the timescale of 5-6 hours (Extended Data Fig. 6n,o and Supplementary Note), neck folding is not driven by tissue buckling due to the opposite cell flows of these two tissues, agreeing with a permissive role of the head and thorax flows. On longer time scale, around 6h post ablation, the deepening speed started to decrease relative to non-ablated control tissue, presumably as a long-term consequence of the tissue stretching observed in the head and thorax cells near the ablation sites (Extended Data Fig. 6n). In the future it will be interesting to explore whether over long time scales surrounding tissue flows contribute to large fold formation during development. Altogether, by exploring large fold formation, we have uncovered how the interplay between in-plane tension and tissue curvature provides a simple, robust, and possibly general mechanism for fold formation. In addition, our work raises novel questions in the control of 3D tissue morphogenesis by the coupled apical and basal tensions, distinct supracellular actomyosin structures and large-scale tissue flows.

So far, in biological systems, curvature has been proposed to play a fundamental role at the subcellular scale in membranes and cytoskeleton dynamics and the role of curvature has been analyzed during cytokinesis, tissue spreading or flows, tumor growth and cell fate specification^37,41, 45–49^. However, the role of curvature in controlling the spatiotemporal dynamics of folding has remained uncharacterized despite theoretical work^50,51^. Our work also extends the study of supracellular actomyosin structures in tissue development and repair^30,37,39,41, 52–55^ by defining how their curvature imposed by animal geometry impinges on the spatiotemporal dynamics of out-of-plane deformations. So far, buckling instabilities driven by tissue compression have been shown on several examples to explain large fold formation, with regional differences in growth or geometry defining their position^25–27,56,57^. Our study exemplifies how in-plane tension combined with tissue compartment geometry induces a large fold at a genetically defined position independently of mechanical instabilities. It further suggests that simple feedback between local tissue geometry and folding velocity can buffer initial spatial variations in tissue geometry. Our findings therefore complement previous results linking folding robustness to the exquisite molecular control of actomyosin dynamics or organization^21,58,59^. Collectively our work highlights a simple design principle driving fold formation and more generally calls for a better understanding of the role of tissue geometry in morphogenesis during development.

## Acknowledgments

We thank Y. Hong, L. Johnston, R. Karess, T. Kaufman, T. Lecuit, V. Mirouse, J. Zallen, the Bloomington, Vienna, Harvard Medical School Stock Centers and the Developmental Studies Hybridoma Bank for reagents; the PICT-IBiSA@BDD imaging facility (ANR-10-INBS-04); A. Bardin, V. Cachoux, F. Gallet, E. Hannezo, P. Léopold, J.-L. Maître, F. di Pietro, D. Pinheiro for comments, O. Renaud, O. Leroy for help with ablation, A. Dauphin for laser power measurements, A. Maugarny-Calès for advice on injections, C. Roffay, S. Rigaud and V. Cachoux for inputs regarding image analyses or visualization. This work was supported by Institut Curie, CNRS, INSERM, ANR-Migrafolds, ERC Advanced (340784), ARC (SL220130607097), ANR Labex DEEP (11-LBX-0044, ANR-10-IDEX-0001-02). A. V. and L. A. acknowledge FRM (FDT201805005805) and ARC (PDF20181208399) fellowships, respectively.

## Author contributions

A.V., L.A., F.G., F.B. and Y.B. designed the project. I.G., A.J. produced reagents. A.V., L.A. and F.B. developed experimental methods and performed experiments. A.V., F.G. developed data analysis and modeling. A.V. developed quantification scripts. A.V., L.A., F.B. analyzed the data. A.V., L.A., F.G., F.B. and Y.B. wrote the manuscript.

## Competing interests

The authors declare no competing financial interests.

## Methods

### Fly stocks and genetics

Supplementary Table 1 lists *Drosophila melanogaster* stocks used and associated references. Loss-of-function and gain-of-function experiments were carried out using the Gal4/UAS system utilizing the temperature sensitive Gal80^ts^^60,61^, except for the *Dfd>Dfd^RNAi^* and *Dfd>Dg^RNAi^* experiments. To tune the transcriptional activity of the *Dfd-GAL4* or *Act-GAL4* drivers, animals were raised at 18°C and incubated at 29°C for 48h prior to imaging or for 8h in the case of *Dfd>Diap1*. As in any temperature shift experiment using the Gal4 system, the magnitude of the loss-of-function depends on the duration of RNAi induction. The observed phenotypes might therefore be stronger at later time point of development. For clonal loss-of-function analyses the FLP/FRT flip-out was employed^62,63^. The *Dfd>Dfd^RNAi^* and *Dfd>Dg^RNAi^* experiments were performed at 25°C. In this condition, a weak Dfd staining can be observed (Extended Data Fig. 3a). Increasing the temperature to 29°C causes early pupal death (or head eversion failure and disc fusion defects) in most of the animals (>50%), precluding analyses of Dfd function in such condition. For clonal loss-of-function analyses larvae were raised at 25°C, heat-shocked at 37°C for 12min (*Tollo^RNAi^*) or 45min (*Dfd^RNAi^*) in the 3^rd^ instar larval stage, and then kept at 29°C for 48h (*Tollo^RNAi^*) or 72h (*Dfd^RNAi^*) before analyses.

### Generation of Dfd-GAL4 and Dg::GFP

The *Dfd-GAL4* and *Dg::GFP* alleles were generated by CRISPR/Cas9-mediated homologous recombination at their endogenous loci, using the vas-Cas9 line^64^. To generate the *Dfd-GAL4* and *Dg::GFP* alleles by CRISPR/Cas9-mediated homologous recombination, guide RNAs were cloned into the *pCFD5:U6:3-t::gRNA* vector^65^. For the *Dfd-GAL4* allele the following guide RNAs were used: 5’- AGA AAA GAG CTC ATG ACG GAA GG-3’ and 5’- CAT GAG AAA AGA GCT CAT GAC GG- 3’; while for the *Dg::GFP* allele we used the following ones: 5’-TGG TTT GCG GTA GGA GGG CGT GG-3’ and 5’-TGG CGG TGG TTT GCG GTA GGA GG-3’. Homology sequences were cloned into homologous recombination vectors harboring a hs-mini-white cassette flanked by two loxP sites^66,67^ and a N-terminal GFP sequence for GFP tagging of Dg or an N-terminal GAL4 sequence to insert GAL4 downstream of the *Dfd* promoter. The vector used for GFP tagging has been described in^66^. The vector used to insert GAL4 was generated by replacing the mKate2 sequence by a GAL4 sequence in the mKate2 tagging vector^66^ using the following primers: 5’-ATG AAG CTA CTG TCT TCT ATC GAA CAA GC-3’ and 5’-ATA GCA TAC ATT ATA CGA AGT TAT GGA TCC TTA CTC TTT TTT TGG GTT TGG TGG GGT ATC-3’. To generate the *Dfd-GAL4* line, the two homologous regions (HR1 and HR2) flanking the site of CRISPR/Cas9 cuts were cloned using the following primers: (HR1) 5’-CCC GGG CTA ATT ATG GGG TGT CGC CCT TCG CGA GGG TAG GTA AGT AGG TGT GC-3’ and 5’-TGC TTG TTC GAT AGA AGA CAG TAG CTT CAT CAT GAC GGA AGG TCT GTT GGA TCG-3’; (HR2) 5’-AGT TCG GGG TCC AGC GGT TCT TCA GGC AGT AGC TCT TTT CTC ATG GGT TAT CCG C-3’ and 5’-ATT TTG TGT CGC CCT TGA ACT CGA TTG ACG CTC TTC GAC TAT GCG GCA TAA CAA TCT TGT GGG C-3’. The following HR1 and HR2 were cloned in the GFP tagging vector for Dg::GFP: (HR1) 5’-CTA ATT ATG GGG TGT CGC CCT TCG GGT CTC TAG TTG AAC GAA GAG TTC TAT GGC ATT CCG-3’ and 5’-GAA CTG CCT GAA GAA CCG CTG GAC CCC GAA CTG GAG GGC GTG GCT GGC GAC TTG-3’; (HR2) 5’-CTC CGG AAG TGG TAG CTC AGG GTC TAG TGG ATA CCG CAA ACC ACC GCC ATA TG-3’ and 5’-GCC CTT GAA CTC GAT TGA CGC TCT TCG TCC GAA TTA TCC AAA GGG GAG CTT GTG-3’. Cloning was performed using SLIC^68^. All embryo transgenesis injections were performed by Bestgene and transgenes were confirmed by sequencing. Detailed plasmid maps and their DNA sequences are available upon request.

### Pupa and adult mounting and imaging

#### Time-lapse live imaging and tissue curvature modulation

Pupae were collected at 0 hAPF or at head eversion (12 hAPF). Pupa mountings for live-imaging were adapted from^66,69^ to reproducibly ensure (i) medial flattening, (ii) lateral flattening or (iii) no flattening of the dorsal neck tissue. At approximately 14-17 hAPF, the dorsal part of the head and thorax pupal case was removed using dissection Vannas spring scissors and Dumont #5 tweezers (Fine Science Tools). *For medial flattening*: Pupae were then adhered ventrally on a slide using double-sided tape (Scotch 3M) and the pupae heads were elevated by placing 6 layers of double-sided tape below them. Two spacers respectively made of six and five 18×18mm #1 (0.13-0.16mm) coverslips (Thermo Scientific, Menzel-Glaser) were placed posteriorly and anteriorly to the pupae, respectively, close to the edges of the slide. The spacers supported the top 24×40mm #1 (0.13-0.16mm) coverslip (Knittel Glass) placed over the pupae and covered with a fine layer of mineral oil 10 S (VWR International). The top coverslip was glued to the spacers using nail polish. This resulted in a flattening of a tissue region centered on the midline. *For lateral flattening*: the mounting was similar except that the pupae were adhered ventrally slightly sideways to yield a lateral flattening upon gluing the top coverslip on the two spacers. *Without coverslip flattening:* to avoid flattening of the dorsal region upon mounting of the top coverslip, the spacers were made respectively of eight and seven 18×18mm coverslips to prevent a direct contact between the coverslip and the pupae. To allow imaging in this case, a thicker layer of mineral oil was spread over the coverslip before placing it on top of the pupae. By gently moving the coverslip when placing it in contact with the two spacers, a contact zone between the mineral oil and the presumptive neck region could be created. Care was taken to restrict immersion into mineral oil to only the most dorsal part of the pupae to prevent the oil from covering the lateral spiracles and disrupting breathing.

Samples were imaged at 25°C or 29°C with an inverted confocal spinning disk microscope from Nikon or Carl Zeiss using 40x/1.4 OIL DIC H/N2 PL FLUOR, 60x/1.4 OIL PL APO, 63x/1.4 OIL DICII PL APO or 100x/1.4 OIL DIC N2 PL APO VC objectives and a sCMOS camera (Orca Flash4, Hamamatsu). For comparison of live-imaging quality with and without coverslip, 18 hAPF Ecad::3xGFP expressing pupae were sequentially imaged using a 40x/0.6 DRY CFI S Plan Fluor ELWD objective without oil and subsequently a 40x/1.4 OIL DIC H/N2 PL FLUOR objective with oil under identical imaging parameters.

To record neck invagination, 14-17 hAPF old Ecad::3xGFP or ubi-Ecad::GFP expressing pupae were imaged until approximately 24 hAPF every 15min, using 60 step *z*-stacks with 1µm spacing. Autofocus detection (Metamorph software) was used to keep the focus on the tissue throughout the time of imaging.

#### Pupal tissue fixation, staining and imaging

Pupae were dissected, fixed and stained as previously described^70^. Primary antibodies used: mouse anti-Antp (DSHB #8C11, 1:20 dilution), rabbit anti-Dfd (1:100 dilution, gift from T. Kaufman). Secondary antibodies used: donkey anti-rabbit Cy5 (Interchim), donkey anti-mouse Cy3 (Interchim). Phalloidin Abberior STAR RED (Sigma, 1:2000) or Phalloidin Alexa Fluor 647 (ThermoFisher, 1:1000) were used to label F-actin. Fixed tissues were mounted in 80% Glycerol (Sigma) in 1xPBS supplemented with 2% N-propyl gallate (Sigma) and imaged using an inverted confocal spinning disk microscope from Nikon or Carl Zeiss using either 40x/1.4 OIL DIC H/N2 PL FLUOR, 60x/1.4 OIL PL APO, 63x/1.4 OIL DICII PL APO or 100x/1.4 OIL DIC N2 PL APO VC objectives and sCMOS camera (Orca Flash4, Hamamatsu).

#### Ex vivo imaging of the basal surface of the pupal dorsal epidermis

To better visualize the basal surface of neck epidermis, an *ex vivo* live or fixed imaging approach was adapted from^71^. Upon removal of the pupal case, pupae were glued on their dorsal side to a 20 mm glass bottom microwell dish (MatTek Life Sciences) using heptane glue^72^. They were then dissected in 1xPBS as previously described^73^ and imaged using an inverted confocal spinning disk microscope from Nikon or Carl Zeiss using either 40x/1.4 OIL DIC H/N2 PL FLUOR, 60x/1.4 OIL PL APO or 100x/1.4 OIL DIC N2 PL APO VC objectives and sCMOS camera (Orca Flash4, Hamamatsu). In case of imaging fixed samples, dissected tissues were fixed and stained as described above.

#### Injections

22h hAPF pupae expressing Ecad::3xGFP were glued on their dorsal side to a 20mm glass bottom microwell dish (MatTek Life Sciences) using heptane glue^72^ and imaged at 10min intervals (80 *z*-stacks with 1µm spacing) for 1.5-2h using an inverted confocal spinning disk microscope from Carl Zeiss using a 40x/1.4 OIL DIC H/N2 PL FLUOR objective. After having recorded the pre-injection neck invagination dynamics, the dish was removed from the microscope and pupae were manually injected under a stereo microscope (Carl Zeiss), using a NanojectII system (Drummond Scientific), either with 27.6nL H_2_O or 50mM Y-27632 (Tocris) dissolved in H_2_O in the abdomen. Upon injection pupae were imaged for an additional 1.5-2h to record the post-injection invagination dynamics.

#### Whole animal imaging

Whole adult flies and pupae were imaged in glycerol:ethanol (4:1) using a Carl Zeiss Stereo Discovery V20 fluorescent microscope equipped with an Axiocam camera using a PlanApoS 1.0x FWD 60mm objective.

### Laser ablations

Laser ablation can be used as a probe, to measure the existing tension, usually by severing the apical, lateral or basal cortex domain of the cells: tissue recoil velocity immediately after ablation is considered to be a reliable proxy of the tension which existed in the tissue immediately before ablation^33,42^. We performed such probe ablation using a multi-photon laser. Laser ablation can also be used to sever the tissue across is whole thickness: it artificially creates a physical opening within the tissue, thus mechanically isolating a piece of tissue from another piece and providing a stress-free boundary. We performed such ablations using a UV 355nm laser.

#### Probe laser ablation and quantification of recoil velocity

To measure the apical and basal tissue recoil velocities after ablation, laser ablations were performed using an inverted laser scanning microscope (LSM880 NLO, Carl Zeiss) equipped with a multi-photon Ti::Sapphire laser (Mai Tai HP DeepSee, Spectra Physics). Ablations were performed in 14 to 24 hAPF pupae expressing the utrABD::GFP actin or Ecad::3xGFP AJ markers mounted with medial coverslip flattening and imaged in single-photon bidirectional scan mode every 1s for 30s with a 40x/1.3 OIL DICII PL APO (UV) VIS-IR (420762-9800) objective. A region of interest (ROI) corresponding to a 34.5×17.25µm rectangle along the AP axis and centered on the medial or lateral neck region was ablated as previously described^69,74,75^. Apical and basal ablations were sequentially performed in the same pupa. The initial tissue recoil velocity, which is proportional to the stress^33^ was determined between *t*=1s and *t*=7s after ablation. No differences in the recoil velocities were found upon changing the order of sequential ablations of the apical and basal domains (Extended Data Fig. 6i,j). To measure AP and ML tension simultaneously, either basally or apically, a small 8.6×8.6µm circular ROI was ablated in 22 hAPF old pupae and imaged every 0.25s. The initial recoil velocity was measured between 0.25s and 1.25s following ablation. Note that recoil velocities after ablation indicate tensions only up to a prefactor (which is typically a dissipation factor). They can be considered as reasonable proxies of tensions^33,42^, while absolute tension values are not measured.

The same microscope set-up was used to measure recoil velocity along the apical-basal axis after lateral ablation. Ablations were performed in 21-22 hAPF Ecad::3xGFP pupae. Before ablation, a 21×21µm stack (50 to 60 *z*-plans, 0.5µm apart) centered on the boundary between thorax and neck was acquired to visualize apical AJ Ecad::GFP signal and the weaker basal Ecad::GFP signal. Ablation was then performed in a 21×21µm ROI located at a *z*-plan equidistant from apical and basal tissue surfaces. The 21×21µm stack (50 to 60 *z*-plans, 0.5µm apart) was subsequently acquired 30s after ablation. The lateral membrane recoil velocity was determined by measuring the distance between apical and basal tissue domains before and 30s after ablation. Since this laser power is sufficient to ablate the actomyosin network of the tissue at a deeper position to trigger tissue recoil, we can safely consider that this regime is sufficient to ablate the lateral cortex to estimate recoil velocity.

#### Tissue severing laser ablation to mechanically isolate neck tissue

To mechanically isolate the presumptive neck tissue from head and thorax tissues, ablations were performed on an inverted spinning disk wide homogenizer confocal microscope CSU-W1 (Roper/Carl Zeiss) equipped with an UV 355nm laser ablation module and a sCMOS camera (Orca Flash4, Hamamatsu). Prior to neck deepening (e.g. prior to 17 hAPF) or at 21 hAPF, a head and a thorax tissue ROI of ∼330×50µm flanking the medial dorsal presumptive neck region were ablated. Ablations were performed using a UV 355nm laser with a power of 0.1mW using the 40x/1.4 OIL DIC H/N2 PL FLUOR objective and across 60 imaged *z*-planes (1µm spacing). Ablations were repeated every 3h. To mechanically isolate the medial presumptive neck tissue from lateral neck tissues, a similar set-up was used to ablate a left and a right ROI (∼50×330µm) flanking the medial neck tissue. Using this ablation regime, we verified that we ablate the tissue across it whole thickness (Extended Data Fig. 11f).

### Quantifications of tissue dynamics

#### Spatial and temporal registration of time-lapse movies

Between 14-17 hAPF the presumptive neck-thorax boundary, the most medial and posterior pair of head macrochaetes, the anterior macrochaetes of the scutum and the midline provide reproducible spatial landmarks easily identifiable using cortical markers such as Ecad::3xGFP (Extended Data Fig. 1c). We used their positions before 18 hAPF to register all time-lapse movies in space. Along the ML *x*-axis, the midline position was defined as Position_ML_ = 0%, and the most posterior and medial pair of head macrochaetes was set at Position_ML_ = -100% (left macrochaeta) and Position_ML_ = 100% (right macrochaeta), respectively (Extended Data Fig. 1c). Along the AP *y*-axis, the initial position of the presumptive neck-thorax boundary in a ML box at Position_ML_ = -50% to 50% was defined as Position_AP_ = 0%, and the average AP position of the anterior scutum macrochaetes was defined as Position_AP_= 100%. In all time-lapse movies, the deviation angle of the AP axis from the vertical axis in the *xy* plane, as well as the deviation angle of the AP axis from the horizontal axis in the *xz* plane did not exceed 15° and were corrected for during data analysis.

To register time-lapse movies in time, we measured in each movie the time at which all microchaete precursors located between Position_AP_ = 20% and 50% underwent their last division and set this time at 22 hAPF. This last division measurement provided a timing with a 30min precision (*N*=11 control animals, data not shown). To correct for the difference in developmental time between 25°C and 29°C, a correction factor of 1.27 was applied to data acquired at 29°C to match the 25°C timing^76^. Note that each experimental condition was compared to a control condition acquired at the same temperature, so this correction factor does not impact any of our conclusions.

#### Tracking of apical and basal neck fold front

The apical neck fold front was tracked in 3D in animals expressing Ecad::3xGFP or ubi-Ecad::GFP from confocal time-lapse movies along the ML *x*-axis, the AP *y*-axis and the AB *z*-axis. The boundary between the presumptive neck cells and the thorax cells (coinciding with the apical fold front) was first tracked in time on *z*-maximal projections. At each time point, a ROI of 10-20µm centered along the boundary (Extended Data Fig. 1c) was used to generate a transverse maximal projection in the ML and apical-basal plane (Extended Data Fig. 1d). Any global imaging drift during acquisition was then corrected using fixed particle dusts on the apical ECM using the transverse view. Note that the boundary is roughly straight on *z*-projections, the curvature of the boundary in the ML and AP plane was therefore not taken into consideration. On each transverse section and using either the Ecad::3xGFP or ubi-Ecad::GFP AJ signal, the apical neck fold front was manually segmented as a line by placing a landmark approximately every 20µm. For quantification, each manually segmented fold front position was then interpolated along its length with regular sampling spacing using the *interparc* function of Matlab (https://www.mathworks.com/matlabcentral/fileexchange/34874-interparc). The basal neck fold front was tracked using a similar approach using the most basal Ecad::3xGFP or ubi-Ecad::GFP signal. Tissue thickness was defined as the distance between the Ecad::3xGFP AJ and basal signals in transverse sections.

#### Neck fold deepening

Neck fold deepening was measured by tracking in time of the apical fold front. Position_ML_ was tracked in time by considering that each Position_ML_ along the front linearly moves in time towards a center of convergence (Extended Data Fig. 1e,f). The center position of convergence was estimated as being at the midline Position_ML_ = 0% and the AB position corresponding to the final position of the neck in a posterior view of an adult head, taking into consideration the impact of flattening due to the mounting procedure (Extended Data Fig. 1e,f). To determine neck fold deepening, for each time point *t* and each Position_ML_, neck fold depth was estimated by calculating the distance between the initial position of the neck fold front at 16 hAPF (or at the onset of imaging for the injection experiments) and the corresponding neck fold front Position_ML_ at subsequent time points. For each animal, neck fold depth was then averaged between Position_ML_= -250% and +250%.

#### Fold front curvature and local normal deepening speed

For each time point *t* and at given Position_ML_ (centered on Position_ML_ = 0% and evenly spaced in 10% increments on the tracked apical fold front, Extended Data Fig. 1g,h), local apical neck fold curvature was estimated by fitting a circle on a triplet of points along the tracked apical fold front corresponding to the ML point at which curvature is estimated and the two points located 22.5µm away on either sides of this ML point. Local curvature was defined as the inverse of the radius of the fitted circle. Using the same triplet of points, the local normal deepening speed was calculated as follows. For each time point *t* and each Position_ML_, normal deepening speed was calculated by measuring the distance between the considered Position_ML_ at time *t* and the intersection between local normal and the fold front Position_ML_ at time *t+1*. Normal deepening speed and curvature were smoothed in time using a 1h45min sliding window. For each time point *t* and each Position_ML_, the average and the standard error of the mean (sem) of front curvature and deepening speed were calculated among animals of a given condition. A few ML (less than 2% of the total) positions located on the edges of the tracked fold front were sampled in less than 5 animals (usually corresponding to Position_ML_ < -300% or Position_ML_ > 300% at the beginning of the time-lapse movies) or showed a curvature characterized by a sem higher than 0.002: they were not taken into consideration in subsequent analyses.

#### Transverse section kymograph

Transverse projections of the neck fold front were used to generate a kymograph by plotting at each time point a 4µm wide region of the tissue centered around Position_ML_ = 140%.

#### Tissue velocity measurements

Local tissue velocities were measured either manually by tracking cell displacements over time in Fiji (see an example in Fig. 1b) or by using particle image velocimetry (PIV) on *z*-maximal projections of the Ecad::3xGFP AJ signal using previously published custom Matlab codes (see an example in Extended Data Fig. 6n)^69^.

#### Detection of the apical surface for apical or basal projection

To analyze protein distributions on the apical or basal surface of the neck fold on a confocal *z*-stack, fluorescent signals were projected using a combination of custom Fiji scripts and Matlab codes that automatically determines the apical *z*-map using an apical marker such as the AJ Ecad, F-actin or MyoII markers. The confocal *z*-stack was first preprocessed in Fiji to detect the most apical position of high variance for each pixel as follows: (i) each *z*-section was filtered using the Fiji Subtract Background plugin (rolling ball radius of 2×0.322µm), the resulting *z*-stack was then downscaled by a factor of 10 and a variance filter was applied along the *z* direction; (ii) using the *peakseek* Matlab function (https://www.mathworks.com/matlabcentral/fileexchange/34874-interparc), the *z*-position of each pixel was detected to define the most apical position of high variance for each pixel; thus generating a topographic map of the apical surface. The topographic map was then up scaled to the original image size to produce the apical *z*-map of the tissue. This apical *z*-map can then be used to project any signal at any apical or basal position.

#### Cell Shape measurements

Measurements of apical and basal AP and ML cell length were performed on PH::GFP *z*-stack images (150 slices at 0.5µm spacing and 0.065µm *x-y* resolution) captured at 16 and 24 hAPF in Fiji by measuring the longest basal and apical cell widths on sagittal and transverse slices centered around the midline and neck regions (∼8 cells wide), respectively. A region of the head-neck-thorax tissues at 16 and 24 hAPF was segmented in 3D by manually tracing the cell outlines in each *z*-slice in Fiji and processed using custom Fiji and Matlab codes. To characterize head, neck and thorax apical cell shape anisotropies, randomly selected cells within the medial neck region (within ∼100µm from the midline) were manually segmented on projected z-stack images of Ecad::3xGFP at 13, 14.5, 16, 18 and 20 hAPF. Aspect ratio of traced apical cell shapes were measured using Fiji.

#### Cell delamination event quantification

Within a cell patch (Position_ML_ = -150% to 150% and Position_AP_ = -25% to 25%) delamination events were manually counted between 16 hAPF and 22 hAPF.

### Quantification of F-actin basal organization

The organization of the F-actin network was quantified on dissected and fixed tissue at 21 hAPF and stained for F-actin. Both apical and basal F-actin signals were projected separately (see *Detection of the apical surface for apical or basal projection)*. The basal F-actin signal intensity was then normalized by the average apical one. The basal F-actin fibers were automatically segmented by (i) applying the Subtract Background Fiji plugin (rolling ball radius 3×0.161µm) and (ii) skeletonized them using a threshold value allowing to segment most of them. The segmented F-actin fibers were then processed using the Fiji Analyze Skeleton plugin to remove F-actin fiber “vertices” and keep a linear portion of each F-actin basal fiber. To ensure a meaningful measurement of F-actin fiber orientation, the orientation of the resulting segmented F-actin basal fibers of length larger than 0.8µm were measured with respect to the ML axis (0° and 90° being, respectively, parallel and perpendicular to ML axis). The length of each fiber was estimated by the distance between its two edges. Average basal F-actin fiber orientation was quantified as a weighted average of the F-actin fiber orientation and length in different ROIs: a 57×123µm ROI centered at the presumptive neck-thorax boundary, a 23.5×123µm ROI located at the neck-thorax boundary on the thorax side or a 23.5×123µm ROI located at the neck-thorax boundary on the head side.

### Quantification of Myosin levels as a function of neck curvature

Neck folding was recorded in pupae expressing both Ecad::3xGFP and MyoII::3xmKate2. Curvature along the ML axis was estimated in 21 hAPF old pupae in the same images using the method described in *Fold front curvature and local normal deepening speed*. The intensity of MyoII::3xmKate2 and Ecad::3xGFP signals were estimated for each time point in top projections centered on the apical plane (see *Detection of the apical surface for apical or basal projection*). First, the background was removed by applying the Subtract Background Fiji plugin (rolling ball radius 200×0.322µm), and signals were smoothed in space using a median filter (size 20×0.322µm). MyoII::3xmKate2 and Ecad::3xGFP levels were then measured along the ML axis by isolating a line manually drawn in the middle of neck region. In each animal, the MyoII::3xmKate2 and Ecad::GFP signals along the ML axis were calculated and binned according to curvature (0.001µm^-1^ bins), to calculate an average value for MyoII::3xmKate2 and Ecad::GFP signals for each curvature bin. For each time-point, the MyoII::3xmKate and Ecad::GFP values of each curvature bin were normalized by the average value for all curvatures. ANOVA test was used to determine whether signal intensity varied across the ML axis.

### Image processing for display

Unless otherwise stated, all images and movies for display represent maximal *z*-projections and were subjected to image processing (denoising and contrast enhancement) using Fiji^77^ or Matlab. 3D representations were obtained using either the Imaris software (https://imaris.oxinst.com/), Interactive 3D Surface Plot plugin in Fiji, or Matlab.

### Statistics

Sample sizes vary in each experiment and animal samples were randomly selected within a given genotype for subsequent analyses. Experiments were repeated at least twice. Only animals correctly mounted for microscopy and without developmental delay were included in the analyses. The numbers of analyzed animals, ablations and analyzed cells are indicated in figure legends. For all graphs, each error bar/error area represents the standard error to the mean (sem), except for ablation, basal fiber orientation and delamination for which standard deviation (sd) is shown. In addition, upon averaging among animals of the same genotype, the sd associated with the slopes of linear fits with a zero intercept were extracted from datasets of deepening speed as a function of the product of curvature (*κ*) and initial recoil velocity estimated by laser ablation (*v*). The corresponding errors are *σ_κ_* and *σ_v_*, respectively, and the error *σ_κ.v_* associated with their product was estimated as: 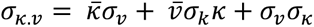 with, respectively, *κ̄* and *v̄* the average curvature and velocity values among animals. Experimental data used for recoil velocity, curvature and deepening speed are given in figure legends. The statistical tests used to assess significance are stated in the figure legends. For comparing deepening dynamics (or the evolution over time of the initial recoil velocity after ablation) between two conditions, a Welch test was performed for each time point, and the associated *p*-value was plotted as a horizontal bar (white: *p*>0.05, striped: *p*<0.05 and plain: *p*<0.01) on top of each graph. Comparisons of two distributions (initial recoil velocities, number of delamination events or orientation of basal fibers) were performed using Welch tests. *t*-tests were performed to analyze whether average changes in cell area, cell AP length and cell ML length were different from 0. ANOVA was used to evaluate the significant differences of MyoII::3xGFP or Ecad::3xmKate2 signal intensities as a function of curvature. Statistical analyses were performed using Matlab or Excel.

### Data and Code availabilities

All data are available upon reasonable requests. Scripts and codes are accessible at: https://github.com/YBellaicheLab/Villidieu_et_al_V1.git

## Supplementary Note

In this Supplementary note, we describe the quantitative analysis and biophysical model that we have used to understand neck folding. Section 1 details the registration and quantification approaches to quantitatively measure neck folding and average it over multiple animals in different mutant contexts. In Section 2 we build upon these quantitative methods, the 3D characterization of cell shape and laser ablation experiments to explore the contributions of apical constriction, apical-basal shortening, cell delamination associated with apoptosis and buckling in the regulation of neck folding. In Section 3, based on the analysis of the distribution of MyoII, the curvature of the Dfd domain and mechanical stress estimation, we then provide a self-contained description of the simple physical consideration leading to the modeling of neck folding. Last, in Section 4 we provide additional considerations regarding the geometrical feedback associated with the stabilization of neck curvature, independently of the initial heterogeneity in curvature.

### 1. Registration and quantification of neck folding and normal deepening speed

In order to quantify neck fold invagination and deepening speed the Ecad::3xGFP signal was imaged in the head-thorax region of a pupa by acquiring *z*-stacks covering a depth of 40µm below the initial tissue plane. Upon *z*-stack projection, the resulting image at 14-17 hAPF was first visualized in top view in order to (i) identify the landmark macrochaetes located in the head and in the thorax necessary for spatial registration along the medial-lateral (Position_ML_) and anterior-posterior axes (Position_AP_) (Extended Data Fig. 1c and see also Methods section for details); and (ii) to track the apical fold front, which is located at the neck-thorax boundary. The *z*-stacks were then resliced (20µm, region indicated by red brackets in Extended Data Fig. 1c) around the location of the apical fold front to obtain transverse views along the ML axis (Extended Data Fig. 1d, blue line defines position of apical fold front). The animal midline position was used as the axis of bilateral symmetry (dashed orange line in Extended Data Fig. 1c-f) and enabled to define coordinates along the ML axis (Position_ML_). The Position_ML_ = 0% can be used as a ML position of the center of convergence of neck invagination since it corresponds to the position of the neck in the adult: its location was estimated using pictures of the back of the adult head on a transverse section (Extended Data Fig. 1e,f). The apical fold front was then tracked in time (Extended Data Fig. 1g). The depth of the apical fold front was then calculated for each Position_ML_ and each time point (Extended Data Fig. 1h). Finally, for each time point, depths were averaged for Position_ML_ between -250% and 250% to obtain the curve of the apical fold front deepening dynamics in each analyzed animal (Extended Data Fig. 1i), then averaged over multiple animals.

### 2. Analyses of the contribution of apical constriction, cell apical-basal shortening, delamination and compressive tissue flows in neck folding

We explored known mechanisms of tissue folding to analyze whether they might explain the observed neck folding: (i) cell apical constriction concomitant with basal relaxation^9–18;^ (ii) apical-basal shortening^19–22;^ (iii) delamination associated with cell apoptosis^23,24^; and (iv) tissue buckling due to compressive stresses^11,25–30^. As shown in Extended Data Fig. 6, our data indicate that these mechanisms do not contribute substantially to neck folding.

i. *Apical constriction concomitant with basal relaxation*: In folds driven by apical constriction, apical cell length reduces in a direction perpendicular to the folding axis, while basal cell length increases; thus leading to cell wedging. This is controlled by distinct apical and basal tension changes in a direction perpendicular to the fold^7^. In contrast, here neck cells did not wedge along the AP axis (i.e. perpendicular to the fold axis) between 16 hAPF and 24 hAPF, but cells reduced their basal sides more than their apical sides (Extended Data Fig. 6a-d). In addition, we found that apical and basal recoil velocities perpendicular to the fold (i.e., AP orientation) are low and similar (Extended Data Fig. 2a). Therefore, the observed cell shape changes and the apical and basal tensions indicate that neck folding is unlikely to be mediated by known apical constriction mechanisms driving folding.
ii. *Apical-basal shortening:* The tissue apical-basal shortening around the fold front corresponded to only a very small fraction of the fold depth (Extended Data Fig. 6e,f). As lateral tension increase has been linked with apical-basal shortening^19–22^, this agrees with the finding that lateral cell tension is too low to be detectable by lateral cell laser ablation (Extended Data Fig. 6g,h). Therefore apical-basal shortening is not a main contributor to neck folding in the pupa.
iii. *Folding driven by delamination associated with apoptosis:* In contrast to apoptosis driven folding^23,24^, here neck cell delamination was not concomitant with a local and transient apical plane bending (Extended Data Fig. 6k). Moreover, reducing apoptosis in the neck by overexpressing the anti-apoptotic *Diap1* gene using a *Dfd-GAL4* driver (*Dfd>Diap1*) did not prevent folding, and if anything, slightly increased its speed (Extended Data Fig. 6l,m). While Dfd has been reported to control segment boundary formation by promoting apoptosis in the *Drosophila* embryo folding^78^, apoptosis does not appear to be a major contributor of neck folding.
iv. *Tissue buckling*: To test the role of putative compressive flows in neck folding, we aimed at abrogating the converging head and thorax tissue flows by laser ablation prior to folding. These laser ablations disrupted the head and thorax flow of cells (Extended Data Fig. 6n), but they did not abrogate the initiation of neck invagination (Extended Data Fig. 6o). Yet, around 6h following ablation, the deepening speed started to decrease relative to non-ablated control tissue, presumably as a long-term consequence of the tissue stretching observed in the head and thorax cells near the ablation sites (Extended Data Fig. 6n,o). We therefore tested whether the head and thorax flows are needed to maintain neck folding by performing similar ablations during folding. When head and thorax tissue flows were abrogated at 21 hAPF, neck folding proceeded normally (Extended Data Fig. 6o). These ablation experiments therefore suggest that head and thorax compressive flows are not required during the initiation and the maintenance phases of neck folding.

Collectively, these experimental findings indicate that the aforementioned mechanisms of tissue folding do not play a major contribution during neck folding.

### 3. Neck folding modeling and energy dissipation

Here we provide a self-contained description of neck folding modeling, as a thin layer which movement is driven by Laplace force and limited by viscous dissipation.

a. *Balance between Laplace and viscous forces*. The magnitude of the Laplace force, per unit length of curve, is the tension *T* multiplied by the curvature “. Since neck folding is associated with large scale tissue flows, we have favored a viscous description of energy dissipation. Inertia being negligible with respect to viscous dissipation, we write that the sum of the Laplace force and of the normal dissipative force is zero; the normal dissipative force being the deepening speed (velocity *v_n_* of the curve along the vector normal to the curve), multiplied by an unknown negative dissipative prefactor, −µ. So, the prediction is that *κT* − µ*v_n_* = 0 or equivalently, that *v_n_* is proportional to *κT* and is directed inwards. Note that this prediction *κT* − µ*v_n_* = 0 is local and involves the tension independently of its cause and of the boundary conditions (whether anchored or periodic).
b. *Thin layer description*. From the theoretical point of view, since the tissue thickness (∼10µm) is smaller than the neck radius (∼300µm) of curvature, the tissue can reasonably be treated as a thin layer. From the experimental point of view, the physical model presented here is a data-driven analysis using the classical equation of Laplace force, of general validity, to perform predictions. Altogether, this model of a line under tension yields correct predictions of neck folding dynamics for both control and mutant conditions in which mechanical tension is altered. We have chosen to treat the apical and basal sides of the tissue separately based on the following experimental and theoretical considerations. We have delineated how genetic regulators, Dfd, Tollo and Dys/Dg determine apical and basal tensions. The apical “tension times curvature” product correlates with the apical tissue velocity, which determines the apical line position versus time; while the basal “tension times curvature” product correlates with the basal velocity which determines the basal line position versus time. The apical quantities are determined with a better signal-to-noise ratio than the basal ones. Both apical and basal velocities are well and separately predicted by the model up to an unknown prefactor, so the absolute values of the velocities are not predicted. The apical and basal prefactors can differ (for instance due to differences in the basal or apical ECM), thus the difference between apical and basal velocities is not predicted. At each time, the difference between apical and basal line positions determines the tissue thickness, which is thus downstream of genetics and velocities, and varies with time. We have quantified this thickness variation with time. Last, our experimental data do not suggest applying the equation of Laplace force to the tissue as a whole to try to predict a « tissue velocity » using the product of a « tissue curvature » with a « tissue tension ». In particular, the velocities, curvature and tensions differ on apical and basal sides, so that defining them at tissue scale might be ambiguous.
c. *Negligible effect from the coverslip on the energy dissipation*. As the neck epithelium harbors an apical ECM (aECM, the cuticle), the coverslip used for live imaging is not in direct contact with the epithelium. In principle, the presence of the coverslip in contact with the aECM could indirectly modulate the friction, the adhesion between the aECM and the epithelium, as well as compress the tissue. The friction of the epithelium on the aECM could induce a force on apical cell surfaces, opposed to the velocity of cells along the coverslip. This, however, would not modify the components of the force and velocity which we observe here, which are perpendicular to the coverslip. Therefore, even if the coverslip-induced flattening changed the friction between the aECM and the epithelium, this would not explain the strong correlation between the deepening speed and the product of curvature and tension. In addition, in principle the coverslip could hinder the cell movements for instance through adhesion. There are strong indications that this is not the case: (i) all the thorax imaging published so far was performed using similar coverslip flattening, yet thorax flow and tissue deformation reproducibly occur at large scale^69,79^; (ii) before 18 hAPF, the neck cells actively move and the neck tissue deforms in plane, hence it is unlikely that cells would be strongly attached (Supplementary Video 1); (iii) in Figs. 4d and 5e, instead of a linear relationship we would have detected a discontinuity (within experimental uncertainty in the curvature measurement, we would have detected at least a significant difference) between the attached part of the tissue, which would have zero deepening velocity, and the non-attached tissue, which is curved and free to invaginate. In summary, the strong linear correlation between the deepening speed and the product of curvature and tension (recoil velocity) can reasonably exclude that the local modulation of neck folding dynamics with coverslip flattening could be explained by the friction or the adhesion between the aECM and the epithelium in the regions flattened by the coverslip.
d. *Negligible effect from the coverslip on other mechanical properties*. The flattening conditions might induce compression on the tissue. Therefore, in principle all the experiments that modulate curvature could also alter concomitantly at least another property of the tissue. In particular, the flattening we impose slightly affects the tension (Extended Data Fig. 11a,b). In principle, it could also affect the mechanical properties of the neck and surrounding tissues, especially through mechanotransduction. Here, the relative change of surface due to the flattening, estimated by comparing the initial surface length before flattening (the arc of circle) with the flattened surface width (the corresponding chord) is of order of 3.7 ± 0.5%. The shear deformation, estimated by the ratio of tissue surface displacement towards the pupa center to the flattened surface width, is of order of 10.6 ± 1.3%. This is well within linear elasticity^80^, “linear” meaning here that the effects are proportional to the cause, and hence likely to remain of order of a few percent too: the modification of mechanical properties is expected to remain moderate. We note that tissue thickness at the midline measured at 18 hAPF has a large variability and is 8.6 ± 1.8µm for flattened animals (*N*=10) vs 7.5 ± 1µm for non-flattened ones (*N*=10); hence the effect of flattening, which is to increase rather to decrease the tissue thickness, is not significant (*p*=0.11). In addition, what we ultimately check is that even if the values of tension, curvature or velocity change, their correlation remains and is compatible with the equation of Laplace force, and thus, with a role of curvature in tuning the spatiotemporal dynamics of neck invagination.

### 4. Robustness and temporal evolution of neck curvature upon flattening

Upon medial flattening of the neck region, the curvature of the midline region gradually increases in time as the neck folds (Extended Data Fig. 11d) and the curvature increase propagates from the more lateral regions to the more medial ones (Extended Data Fig. 11e). Therefore, while the medial neck region is initially flat, over time its curvature becomes like the one observed in the absence of medial flattening. Such stabilization of the neck curvature is a property of curvature-driven movements that robustly yields a homogeneous curvature when shrinking. In fact, consider a curve in which the curved regions move faster than less curved ones. Then, for purely geometrical reasons, when this curve shrinks the curved regions shrink faster, the less curved regions shrink more slowly, and their curvature increases until it equals that of the more curved regions. Hence, the curve shape robustly becomes an arc of a circle (i.e., has a spatially homogenous curvature). Note that in principle another known mechanism could lead to curvature homogenization, namely if the dissipation coefficient µ was anisotropic as is the case when the curve moves within a crystalline lattice^81,82^. Here, because of the experimental quantitative linear agreement of the product of tension and curvature with deepening speed, such anisotropic mechanism is excluded.

---

**Extended Data Fig. 1:**
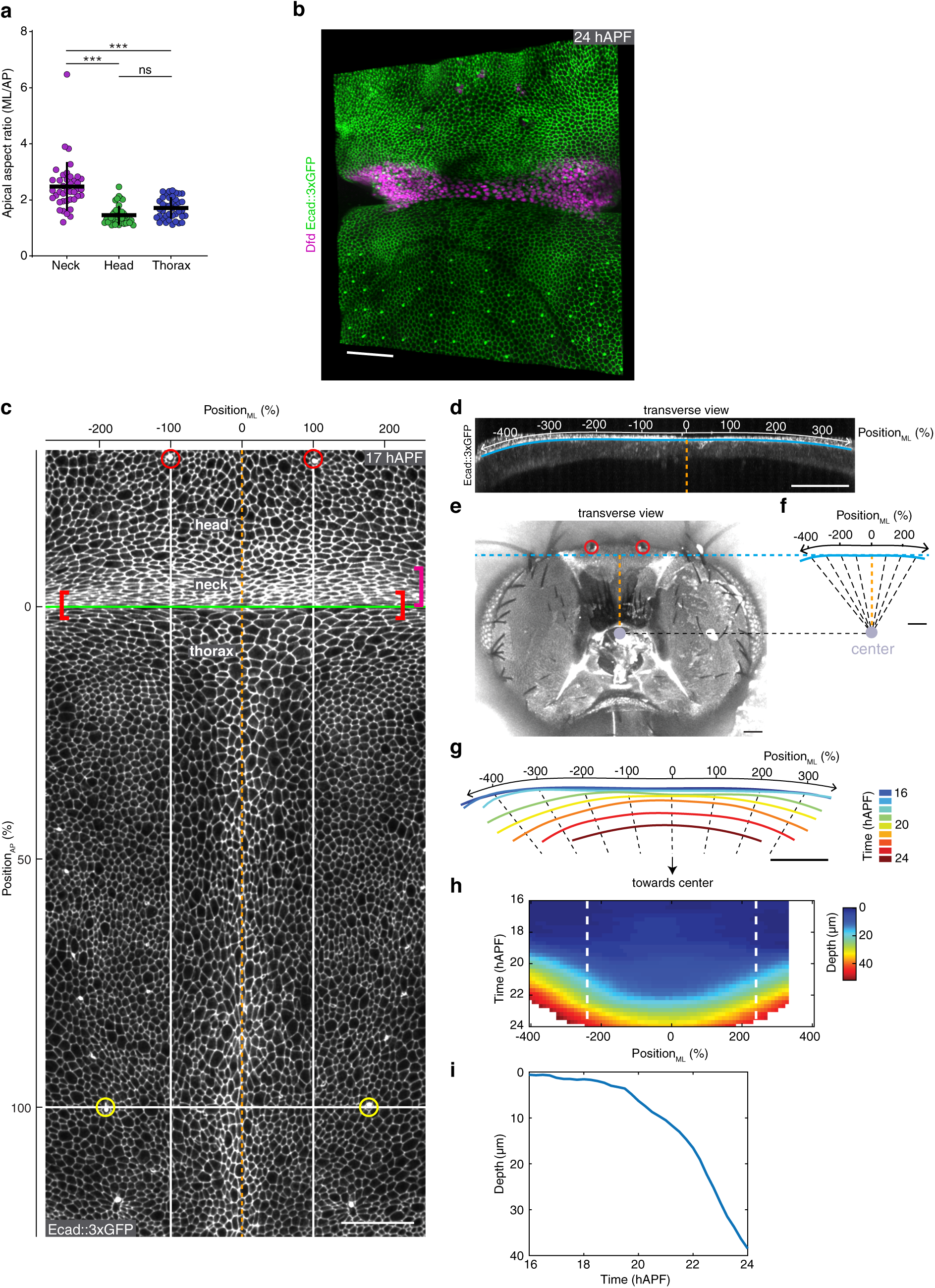
Quantification of neck folding dynamics. **(a)** Graph of the neck, head, and thorax apical cell anisotropy at 13 hAPF (*n=*40 cells from *N=*2 pupae for each tissue region). **(b)** 3D image of Ecad::3xGFP and Dfd localizations in the dorsal neck region at 24 hAPF to illustrate the distribution of Dfd in the lateral region of the neck domain in the most lateral region of the neck region accessible with our imaging setup. The cropped version is shown in Fig. 1c. **(c)** Top view image of the head, neck and thorax regions labelled by Ecad::3xGFP at 16 hAPF. Red and yellow circles, landmark macrochaetae of the head and thorax, respectively, used to define the coordinates along the ML (Position_ML_) and AP (Position_AP_) axes. Green line, neck-thorax boundary. Orange dashed line, midline position. Orange brackets at the neck-thorax interface corresponds to a 20µm box used to track the apical fold front in transverse sections. The magenta bracket indicates the neck region. **(d)** Transverse view at 16 hAPF of the apical fold front region labelled by Ecad::3xGFP. Position_ML_ are indicated. Blue line, apical fold front. Orange dashed line, midline position. **(e)** Transverse view of the back of the adult head. Blue dashed line: estimated flattening applied by the coverslip on the animal during imaging. Grey circle: estimated center of convergence of neck folding. Orange dashed line, axis of bilateral symmetry and distance between midline and center (approximately 200µm). Red circles, landmark macrochaetae of the head. **(f)** Position of apical fold front on transverse view at 16 hAPF, plotted with the estimated center of convergence of neck invagination (grey circle). Orange dashed line, axis of bilateral symmetry and distance between midline and center (approximately 200µm). The black dashed lines indicate the approximate successive positions of points at given Position_ML_ during neck folding. **(g)** Color-coded successive positions of the apical fold front for the animal shown in (d) tracked over developmental time. Black dashed lines represent the estimated successive positions of points at given Position_ML_ during neck folding. The successive positions were determined using the estimated center of convergence (see e,f). **(h)** Graph of the depth of apical fold front as a function of Position_ML_ and developmental time. White dashed lines delimit the +250% and -250% Position_ML_ between which fold depth is averaged at each time point to generate the graph in (i). **(i)** Graph of the average depth of apical fold front (between Position_ML_ = -250% and 250%) of the animal in (d) versus developmental time. Scale bars, 40µm (b), 50µm (c-g).

**Extended Data Fig. 2:**
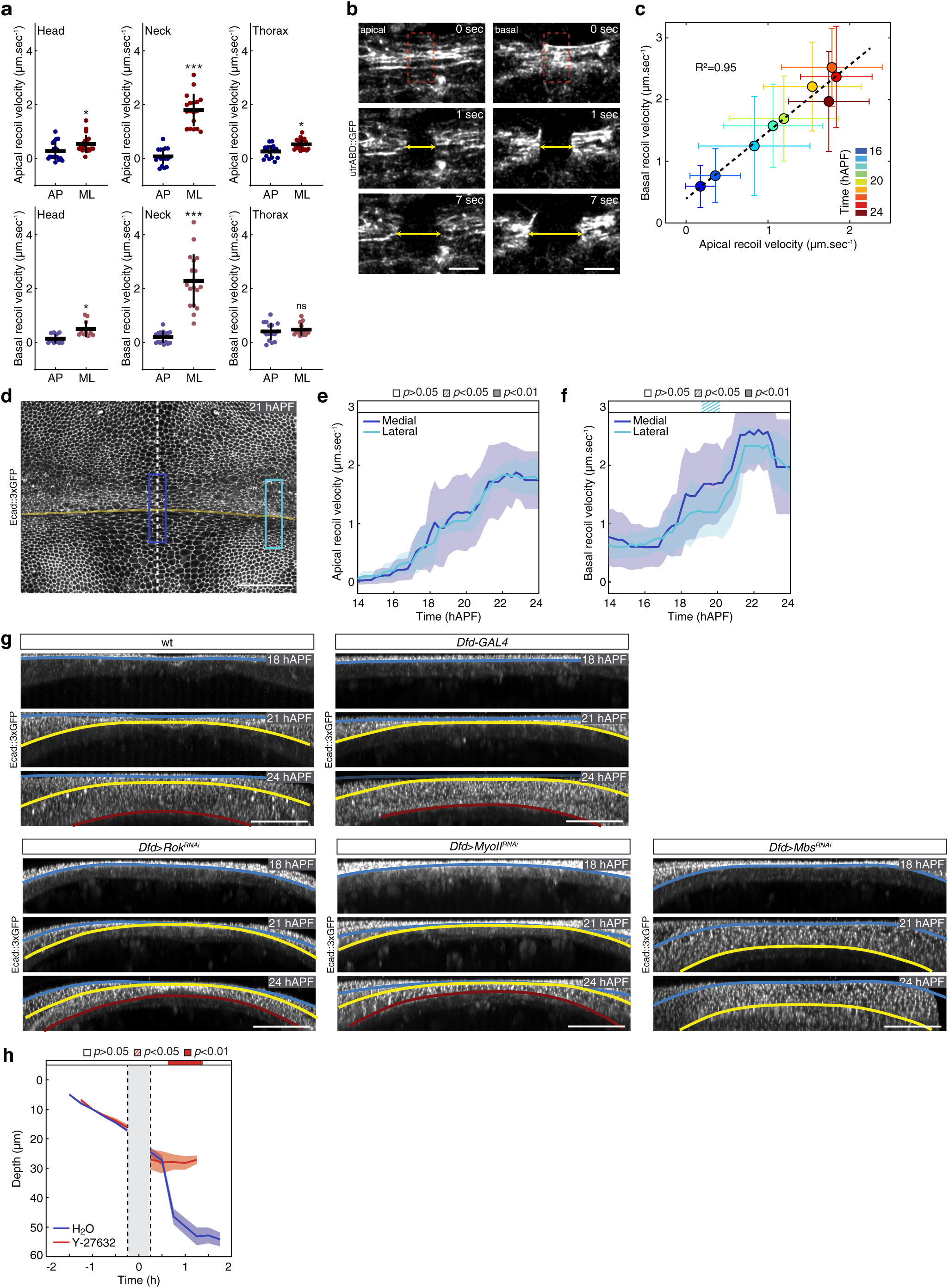
Actin organization, measurements of apical and basal tensions and neck apical folding. **(a)** Graph of the apical (top panels) and basal (bottom panels) initial recoil velocities (mean ± sd) after ablations measured along the AP and ML axes at 22 hAPF in head (apical *N=*18 pupae, basal *N=*10 pupae), neck (apical and basal *N=*18 pupae) and thorax (apical *N=*19 pupae, basal *N=*16 pupae) regions to estimate AP and ML tissue tension. Welch test, ns *p*>0.5, * *p*<0.05, *** *p*<0.001. Circular ablations were used to measure recoil simultaneously along both the AP and the ML axes. As AP tension is negligible all further reported measurements of recoil velocities were performed in a rectangular ROI (b) to analyze ML tissue recoil velocities. **(b)** Top view images of utrABD::GFP actin labelling in the apical and basal neck tissue at 22 hAPF prior to ML ablation (*t*=0s) and after ablation (*t*=7s) in the indicated ROIs (red dashed boxes). Yellow arrows indicate tissue recoil used to measure tissue recoil velocity between *t*=1s and *t*=7s to estimate tissue tension. **(c)** Graph of the apical versus basal ML recoil velocity after laser ablation at different developmental times. Each point is the mean value (± sd) among 127 animals for a given color-coded developmental time. The correlation between the two recoil velocities is indicated (R^2^=0.95). **(d)** Top view image of the neck region labelled by Ecad::3xGPF at 21 hAPF indicating the two regions (medial, dark blue and lateral, light blue) where apical and basal recoil velocities were measured after laser ablation (see e and f). Dashed line, midline position. Yellow line, neck-thorax boundary. **(e)** Graph of the ML apical initial recoil (mean ± sd, averaged with a 2h sliding window) velocities measured upon ablation in the medial (dark blue, *N=*127 pupae) and lateral (light blue, *N=*45 pupae) neck regions as a function of developmental time. Horizontal boxes: *p*-values of Welch tests performed between pupae ablated medially and laterally at successive time points (white *p*>0.05, striped *p*<0.05, solid *p*<0.01). **(f)** Graph of the ML basal initial recoil (mean ± sd, averaged with a 2h sliding window) velocities measured upon ablation in the medial (dark blue, *N=*127 pupae) and lateral (light blue, *N=*45 pupae) neck regions as a function of developmental time. Horizontal boxes: *p*-values of Welch tests performed between pupae ablated medially or laterally at successive time points (white *p*>0.05, striped *p*<0.05, solid *p*<0.01). **(g)** Transverse view time-lapse images of the neck region at 18, 21 and 24 hAPF in wt and *Dfd-GAL4* control conditions, as well as in *Dfd>Rok^RNAi^*, *Dfd>MyoII^RNAi^*, *Dfd>Mbs^RNAi^* representative animals. Tissues are labelled with Ecad::3xGFP. Colored lines outline the position of the apical fold front at successive time points (blue 18 hAPF, yellow 21 hAPF, red 24 hAPF). Note that in the *Dfd>Mbs^RNAi^* the tissue has moved out of the field view at 24 hAPF and the apical fold front line cannot be shown. **(h)** Graph of the apical neck depth (blue, mean ± sem) in pupae mock injected with H_2_O (blue, *N*=6 pupae) and 50mM Y-27632 (red, *N*=8 pupae) as a function of developmental time. Apical neck depth is defined relative to the initial position of the AJ labeled by Ecad::3xGFP at the onset of imaging. The region in grey corresponds to the period during which the injection was performed and time-lapse imaging was stopped (see Methods for details). Horizontal boxes: *p*-values of Welch tests performed between pupae injected with H_2_O or Y-27632 at successive time points (white *p*>0.05, striped *p*<0.05, solid *p*<0.01). Scale bars, 20µm (b), 50µm (d,g).

**Extended Data Fig. 3:**
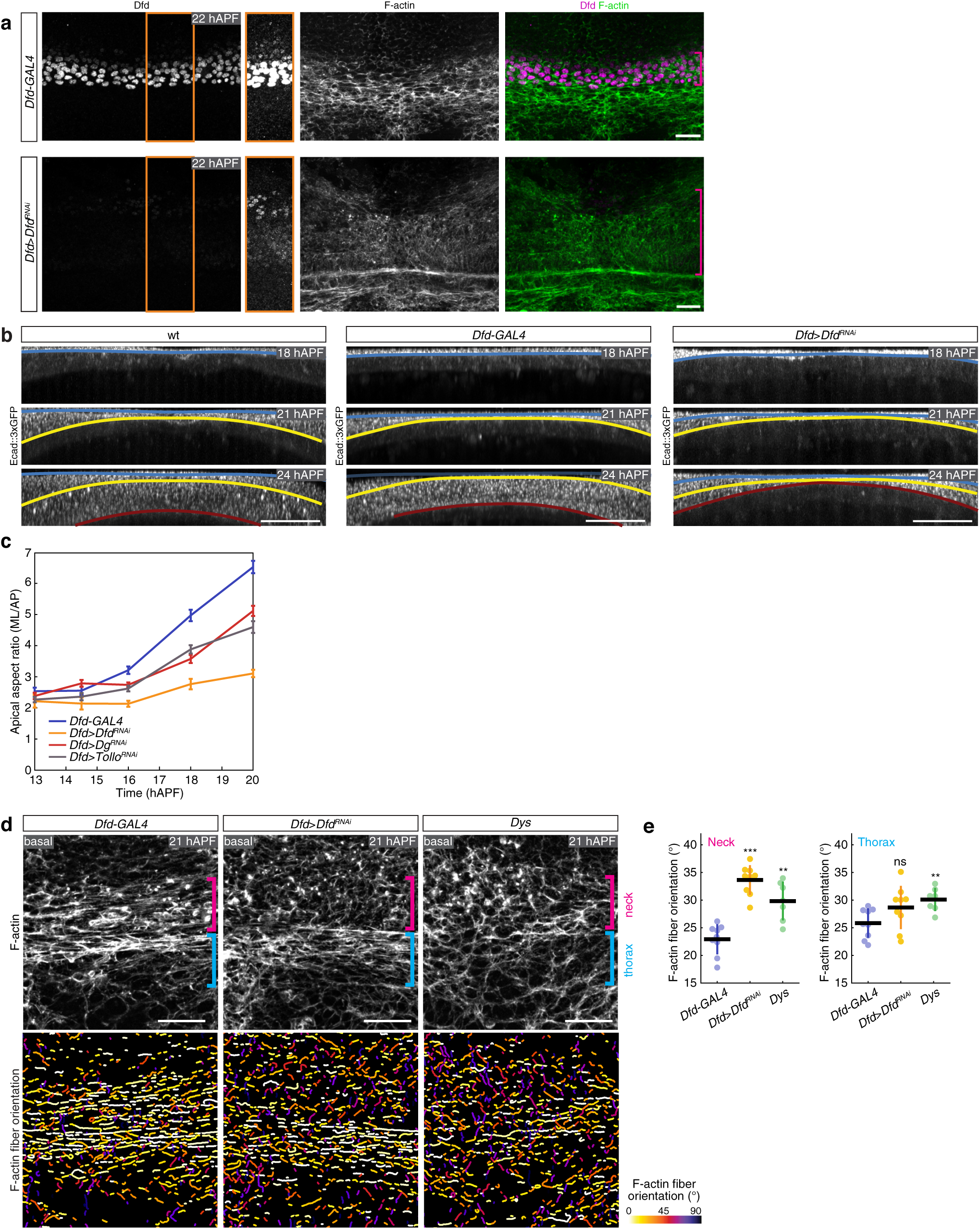
*Dfd*, *Tollo* and *Dys*/*Dg* regulate the assembly of neck apical and basal actomyosin structures and neck folding dynamics. **(a)** Dfd (grey left, magenta right) and F-Actin (grey middle, green rigth) localization in the neck region at 22 hAPF in control *Dfd*>*GAL4* and *Dfd*>*Dfd^RNAi^* mutant tissues. The regions outlined in orange on the left panels are shown next to it with a higher gain. Dfd signal can be detected in the neck in *Dfd*>*Dfd^RNAi^* mutant tissues and the neck region appears enlarged in this mutant condition. In particular, the most anterior domain of the Dfd expression domain does not seem to be affected. Magenta brackets, neck region. **(b)** Transverse view time-lapse images of the neck region at 18, 21 and 24 hAPF in wt and *Dfd-GAL4* control conditions, as well as in a *Dfd>Dfd^RNAi^* representative animal. Tissues are labelled with Ecad::3xGFP. Colored lines outline the position of the apical fold front at successive time points (blue 18 hAPF, yellow 21 hAPF, red 24 hAPF). **(c)** Graph of the neck apical cell anisotropy for *Dfd-GAL4* (*n*=45 cells from *N*=3 pupae for 13 and 14.5 hAPF, *n*=105 cells from *N*=7 pupae for 16, 18, and 20 hAPF), *Dfd>Dfd^RNAi^* (*n*=30 cells from *N*=2 pupae for 13 and 14.5 hAPF, *n*=105 cells from *N*=7 pupae for 16, 18, and 20 hAPF), *Dfd>Dg^RNAi^* (*n*=45 cells from *N*=3 pupae for 13 and 14.5 hAPF, *n*=120 cells from *N*=8 pupae for 16, 18, and 20 hAPF) and *Dfd>Tollo^RNAi^* (*n*=45 cells from *N*=3 pupae for 13 and 14.5 hAPF, *n*=105 cells from *N*=7 pupae for 16, 18, and 20 hAPF) between 13 and 20 hAPF. **(d)** Basal F-actin localization (top) and segmentation of the F-actin fibers color-coded by orientation (bottom) at 21 hAPF in control *Dfd*>*GAL4*, *Dfd*>*Dfd^RNAi^* and *Dys* mutant tissues. Segmented F-actin fibers are colored according to their orientation, ranging from 0° (along ML axis) to 90° (along the AP axis). While F-Actin fiber orientation is affected in *Dfd*>*Dfd^RNAi^* and *Dys* mutant tissues (e), the phalloidin signal level is similar to the ones observed in control *Dfd*>*GAL4* mutant tissues. Blue and magenta brackets indicate the thorax and neck regions respectively where F-actin fiber orientation was analyzed. **(e)** Graph of the F-actin fibers orientation relative the ML axis (mean ± sd) in control *Dfd*-*GAL4* (*N=*9 pupae), *Dfd*>*Dfd^RNAi^* (*N=*9 pupae) and *Dys* (*N=*7 pupae) mutant tissues quantified in the neck (left) or the thorax (right) regions at 21 hAPF. Each F-actin fiber orientation is measured between 0 and 90° relative to the ML axis (i.e., 0° corresponding to a fiber parallel to the ML axis). Each dot is the fiber orientation averaged for one animal. Welch test, ns *p*>0.5, ** *p*<0.05, *** *p*<0.001. Scale bars, 20µm (a,d), 50µm (b).

**Extended Data Fig. 4:**
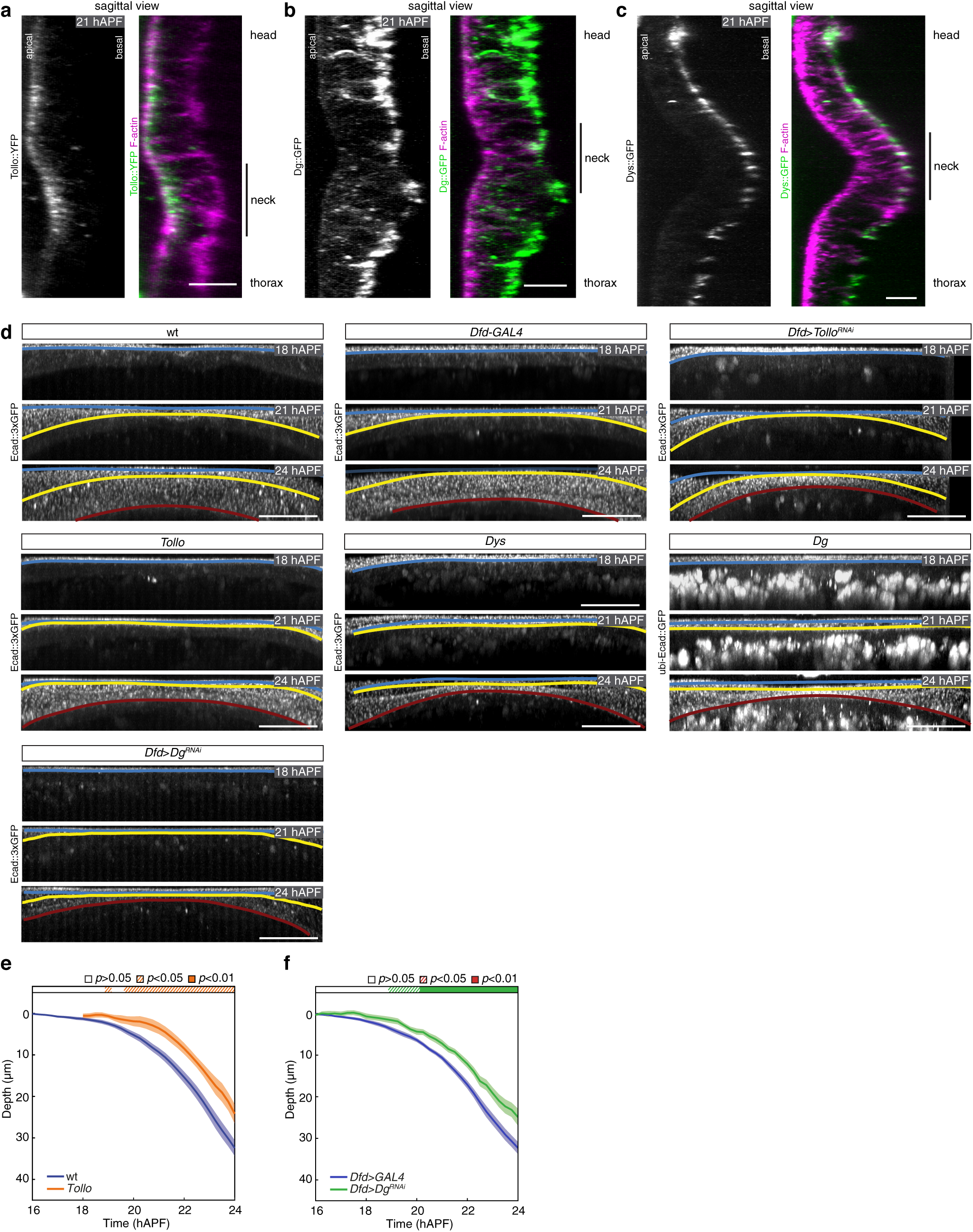
Tollo and Dys/Dg mutant neck folding dynamics. **(a)** Sagittal view of Tollo::YFP (grey left, green right) and F-actin (magenta right) distributions in the neck region at 21 hAPF. **(b)** Sagittal view of Dg::GFP (grey left, green right) and F-actin (magenta right) distributions in the neck region at 21 hAPF. **(c)** Sagittal view of Dys::GFP (grey left, green right) and F-actin (magenta right) distributions in the neck region at 21 hAPF. **(d)** Transverse view time-lapse images of the neck region at 18, 21 and 24 hAPF in wt and *Dfd-GAL4* control conditions, as well as in *Dfd>Tollo^RNAi^*, *Tollo*, *Dys*, *Dg* and *Dfd>Dg^RNAi^* representative animals. Tissues are labelled with Ecad::3xGFP, except for *Dg* which is labelled with ubi-Ecad::GFP. The ubi:Ecad::GFP transgene also promotes Ecad::GFP expression in circulating cells that are present below the tissue. Colored lines outline the position of the apical fold front at successive time points (blue 18 hAPF, yellow 21 hAPF, red 24 hAPF). **(e)** Graph of apical neck depth (mean ± sem) in wt (*N*=10 pupae) and *Tollo* (*N*=7 pupae) mutant tissues as a function of developmental time. Horizontal box: *p*-values of Welch tests performed between the wt control and *Tollo* mutant condition at successive time points (white *p*>0.05, striped *p*<0.05, solid *p*<0.01). **(f)** Graph of apical neck depth (mean ± sem) in *Dfd-GAL4* (*N*=9 pupae) and *Dfd>Dg^RNAi^* (*N*=9 pupae) mutant tissues as a function of developmental time. Horizontal box: *p*-values of Welch tests performed between the *Dfd-GAL4* control and *Dfd>Dg^RNAi^* mutant condition at successive time points (white *p*>0.05, striped *p*<0.05, solid *p*<0.01). Scale bars, 10µm (a-c), 50µm (d).

**Extended Data Fig. 5:**
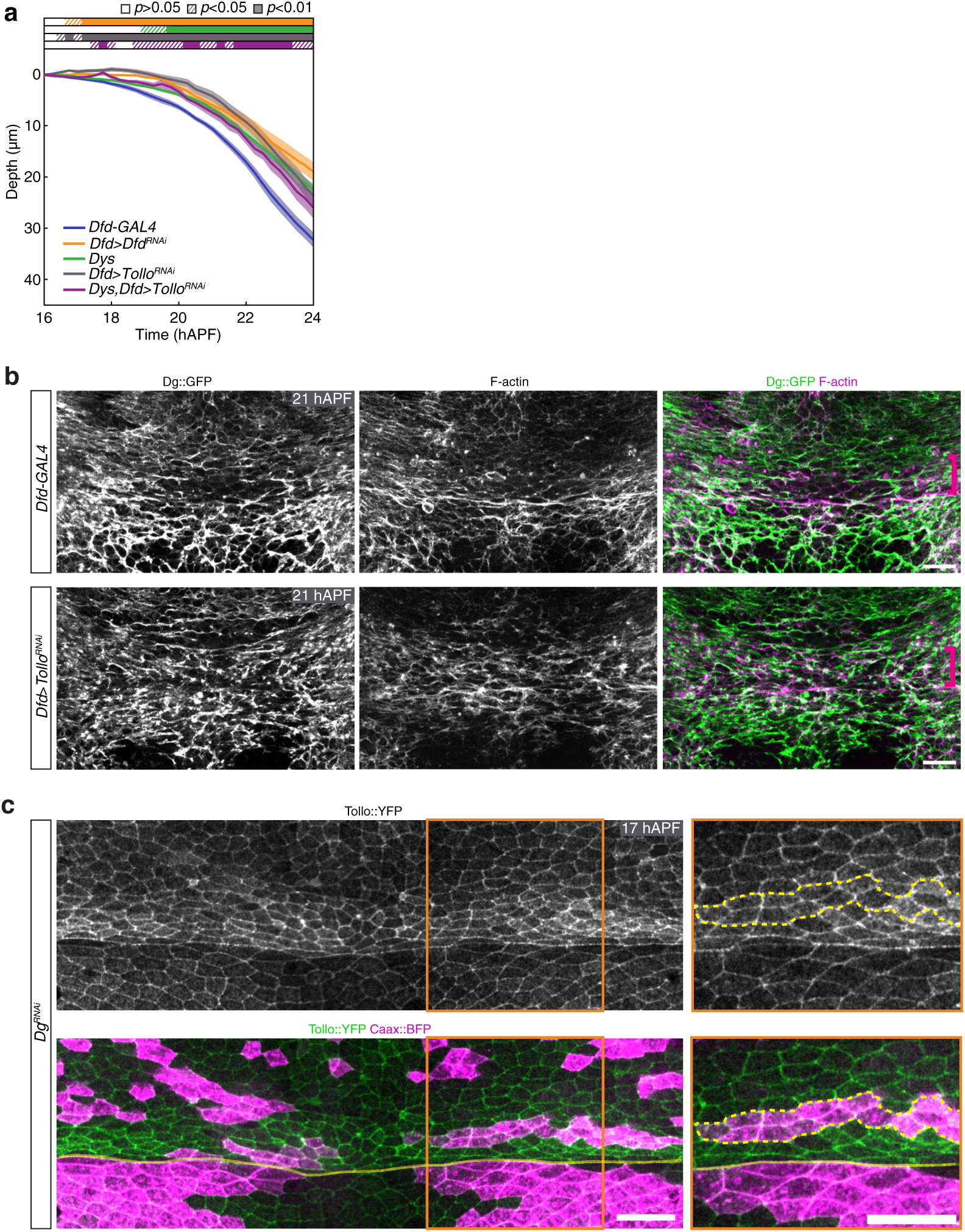
Interplays between Tollo and Dg function during neck folding. **(a)** Graph of apical neck depth (mean ± sem) in *Dfd-GAl4* control (*N=*9 pupae), *Dfd>Dfd^RNAi^* (*N=*8 pupae), *Dys* (*N=*12 pupae), *Dfd>Tollo^RNAi^* (*N=*11 pupae) and *Dys,Dfd>Tollo^RNAi^* (*N=*8 pupae) mutant tissues as a function of developmental time. Horizontal boxes: *p*-values of Welch tests performed between the *Dfd-GAl4* control and *Dfd>Dfd^RNAi^* (orange) or *Dfd>Tollo^RNAi^* (dark grey) or *Dys,Dfd>Tollo^RNAi^* (purple) as well as between wt control and *Dys* (green) at successive time points (white *p*>0.05, striped *p*<0.05, solid *p*<0.01). **(b)** Basal Dg::GFP (grey left, green right) and F-actin (grey middle, green right) distributions in control *Dfd*>*GAL4* (top) and *Dfd*>*Tollo^RNAi^* (bottom) mutant tissues at 21 hAPF. Magenta brackets, neck region. **(c)** Apical Tollo::YFP (grey top, green bottom) in *Dg^RNAi^* clones, marked by Caax::BFP accumulation (magenta bottom right) at 21 hAPF. Yellow dashed line, outlines of *Dg^RNAi^* clones. A close up of the region outlined in orange is shown next to the panel. Yellow line, neck-thorax boundary. Scale bars, 20µm.

**Extended Data Fig. 6:**
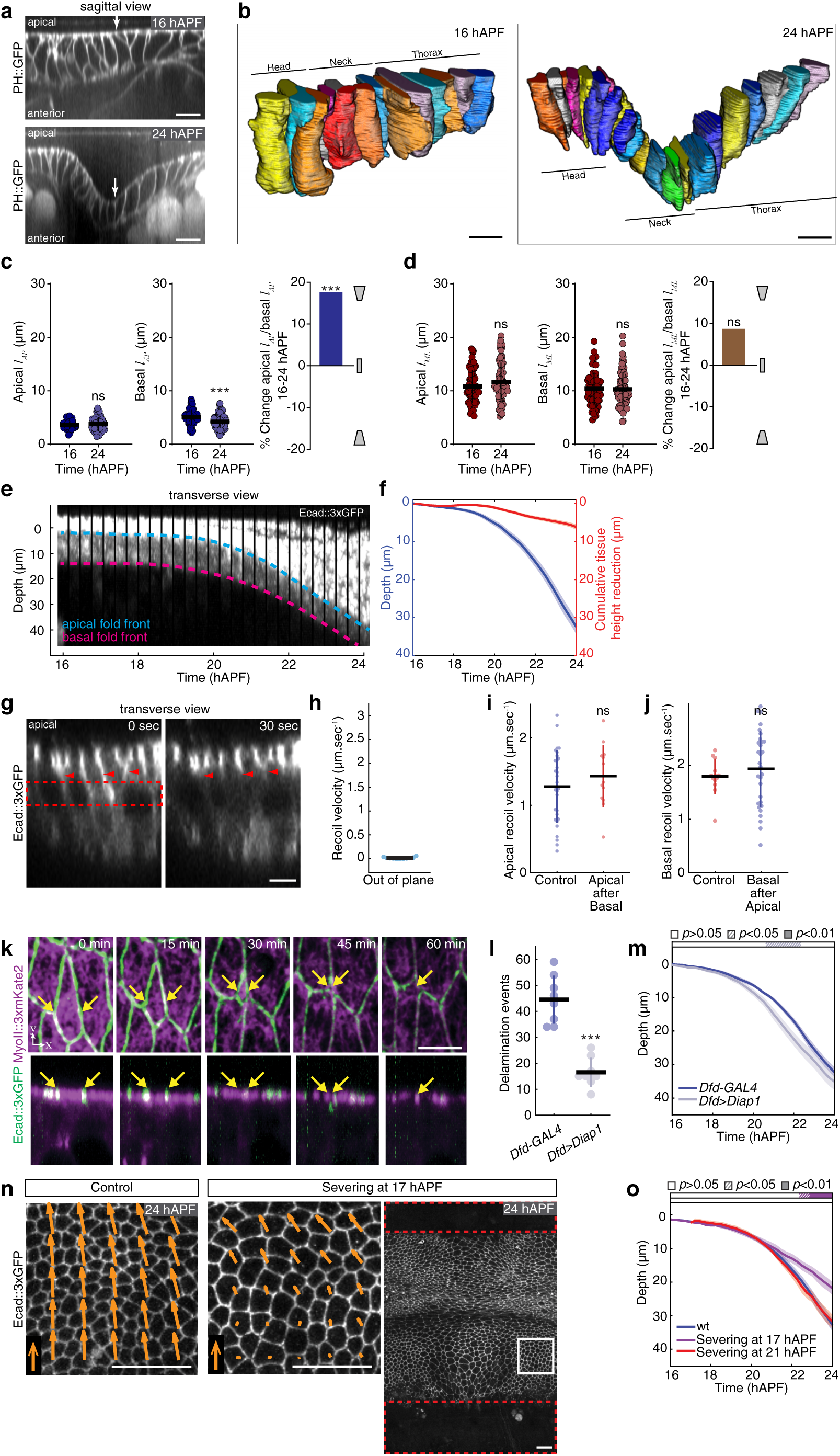
Analyses of the contribution of apical constriction, cell apical-basal shortening, delamination and compressive tissue flows in neck folding (see also Supplementary Note). **(a)** Sagittal view images of the PH::GFP labelled cell membranes in the neck region at 16 and 24 hAPF. Arrows indicate neck-thorax interface. **(b)** 3D views of a row of cells in the head, neck and thorax regions centered around the midline at 16 and 24 hAPF. **(c)** Graphs of the apical (left) and basal (middle) cell length (mean ± sd) along the AP axis (*l*_AP_) at 16 hAPF and 24 hAPF and of the apical-basal cell shape changes along the AP axis between 16 and 24 hAPF (right) measured using the PH::GFP marker. The apical-basal cell shape change along the AP axis was measured as the ratio between the apical and basal cell length variation along this axis. For each time point 75 cells were measured from *N*=3 pupae. *t*-test: ns *p*>0.05, *** *p*<0.001. **(d)** Graphs of the apical (left) and basal (middle) cell length (mean ± sd) along the ML axis (*l*_ML_) at 16 hAPF and 22 hAPF and of the apical-basal cell shape changes along the ML axis between 16 and 24 hAPF (right) measured using the PH::GFP marker. The apical-basal cell shape change along the ML axis was measured as the ratio between the apical and basal cell length variation along this axis. For each time point 75 cells were measured from *N*=3 pupae. *t*-test, ns *p*>0.05. **(e)** Transverse view kymograph of the neck region labelled by Ecad:3xGPF from 16 to 24 hAPF. Blue and pink dashed lines indicate the successive apical (blue) and basal (pink) fold front positions of the neck tissue, respectively. **(f)** Graph of apical neck depth (blue, mean ± sem) and of cumulative tissue height reduction (red, mean ± sem) as a function of developmental time (*N*=10 pupae). Apical neck depth is defined relative to the initial position of the AJ labeled by Ecad::3xGFP. Tissue height is defined by the difference between the apical and basal Ecad::3xGFP signal (see e). **(g)** Images of apical-basal Ecad::3xGFP distribution on a transverse view in the neck region at 21 hAPF prior to ablation and 30 sec after ablation of the lateral region (red dashed box). Arrowheads, positions of Ecad::3xGFP labelled tissue prior to and after ablation. Since this laser power is sufficient to ablate the actomyosin network of the tissue at a deeper position to trigger tissue recoil, we can safely consider that this regime is sufficient to ablate the lateral cortex to estimate recoil velocity. **(h)** Graph of the out of plane (*N*=11 pupae) recoil velocity (mean ± sd) upon lateral tissue ablation. **(i)** Graph of the ML apical initial recoil velocities in the medial neck region (mean ± sd) without (control) (*N*=30 pupae) and after a preceding basal tissue ablation (*N*=13 pupae). *t*-test, ns *p*>0.05. Ablations were performed at 22 ± 1 hAPF. **(j)** Graph of the ML basal initial recoil velocities in the medial neck region (mean ± sd) without (control) (*N*=13 pupae) and after a preceding apical tissue ablation (*N*=30 pupae). *t*-test, ns *p*>0.05. Ablations were performed at 22 ± 1 hAPF. **(k)** Top (top panels) and lateral view (bottom panels) time-lapse images of Ecad::3xGFP and MyoII::3xmKate2 in a neck cell which is delaminating. Yellow arrows, contours of the delaminating cell. Note the absence of tissue bending in the apical plane during delamination (bottom panels). **(l)** Graph of delamination events (mean ± sd) from 14 hAPF to 21 hAPF in control tissue (*Dfd-GAL4*, *N*=8 pupae) and tissue overexpressing Diap1 (*Dfd>Diap1*, *N*=8 pupae). Delamination events were quantified in an equivalent neck region centered around the midline in all animals. Welch test, *** *p*<0.001. **(m)** Graph of apical neck depth (mean ± sem) in control *Dfd-GAL4* (*N*=9 pupae) and *Dfd>Diap1* (*N*=12 pupae) tissues as a function of developmental time. Horizontal box: *p*-values of Welch tests performed between the *Dfd-GAL4* control and *Dfd>Diap1* mutant condition at successive time points (white *p*>0.05, striped *p*<0.05, solid *p*<0.01). **(n)** Distribution of Ecad::3xGFP at 24 hAPF in control wt (left) and ablated wt pupa (right, dashed red boxes) at 17 hAPF in the head and in the thorax tissue prior to folding onset. White box indicates region shown in center panel. In the left panel an equivalent region is shown in the control tissue. Orange arrows, average flow speed vectors from 21 hAPF to 24 hAPF measured by PIV (5µm.h^-1^, orange arrows in the bottom left). Note that tissue flows are strongly reduced except very close to the invagination position. Also note that in the ablated pupa, the cells are stretched along the AP axis (compare control, left panel with ablated, middle panel). **(o)** Graph of apical neck depth (mean ± sem) in control wt pupae (*N*=10 pupae) and in wt pupae where head and thorax tissue were ablated prior to folding (17 hAPF) (*N*=5 pupae) or ablated at 21 hAPF (*N*=7 pupae) as a function of developmental time. Horizontal boxes: *p*-values of Welch tests performed between experimental condition and the wt control at successive time points color coded according to experimental condition (white *p*>0.05, striped *p*<0.05, solid *p*<0.01). Scale bars, 5µm (g,k), 10µm (a,b), 50µm (n).

**Extended Data Fig. 7:**
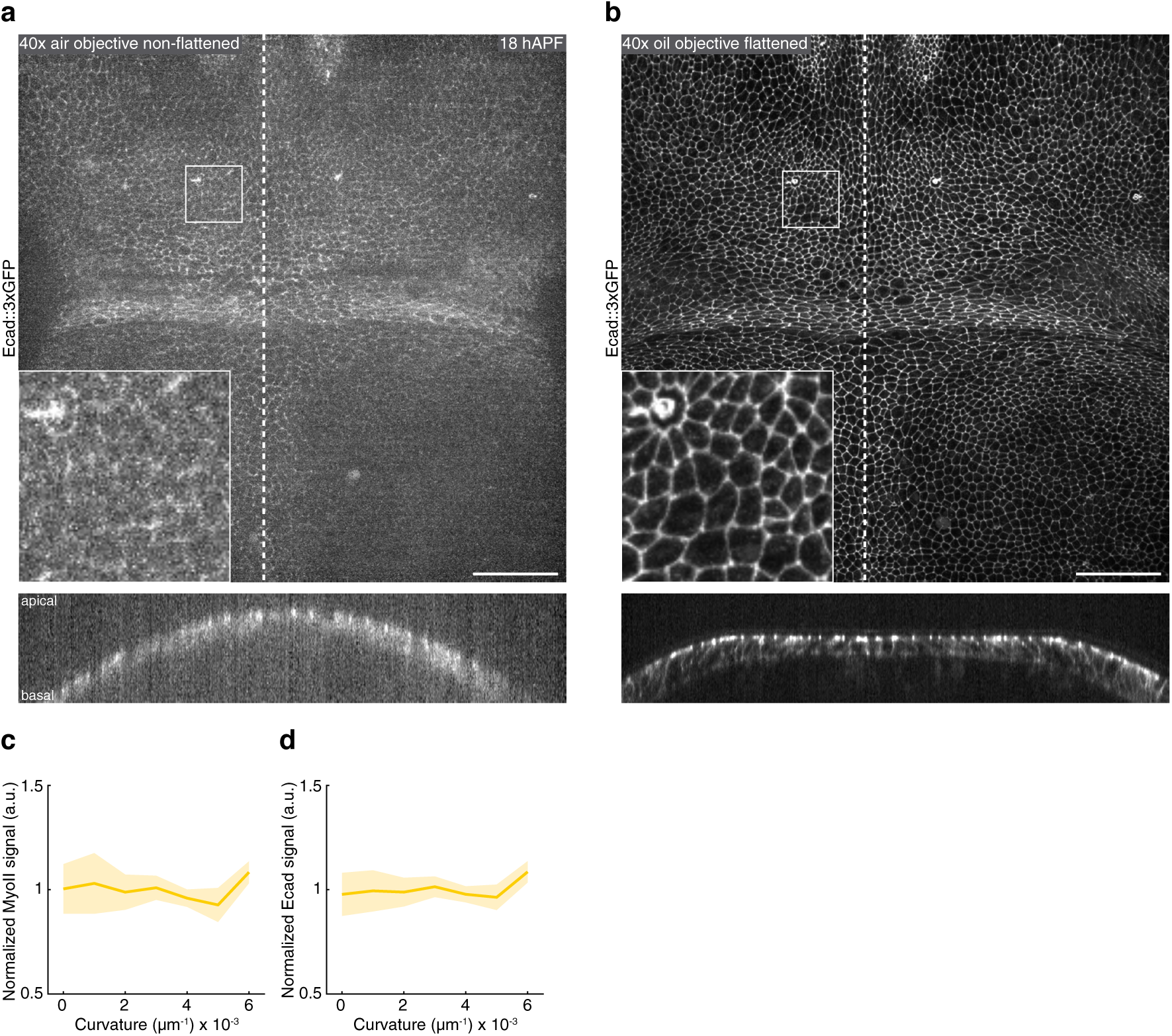
Imaging with and without coverslip as well as Myosin and Ecad distributions as a function of curvature. **(a,b)** Top view (top panels) and transverse view (bottom panels) images of the neck region labelled by Ecad::3xGPF at 18 hAPF of the same pupa imaged without coverslip using an air objective (a) and with a coverslip that locally flattened the tissue using an oil objective (b). Image acquisition settings (laser power and exposure time) are identical in the two conditions. As expected, the image quality is higher using an oil objective and a coverslip. Dashed line, midline position. Inset: enlargement of the square. **(c,d)** Graphs of the normalized MyoII (c) and Ecad (d) signal (± sem) in the neck region along the ML axis as a function of the local tissue curvature in 21 hAPF pupae imaged with medial coverslip flattening (*N=*10 pupae). While the curvature is spatially heterogeneous, the MyoII and Ecad signals remain homogeneous along the ML axis. ANOVA, *p*=0.9. Scale bars, 50µm.

**Extended Data Fig. 8:**
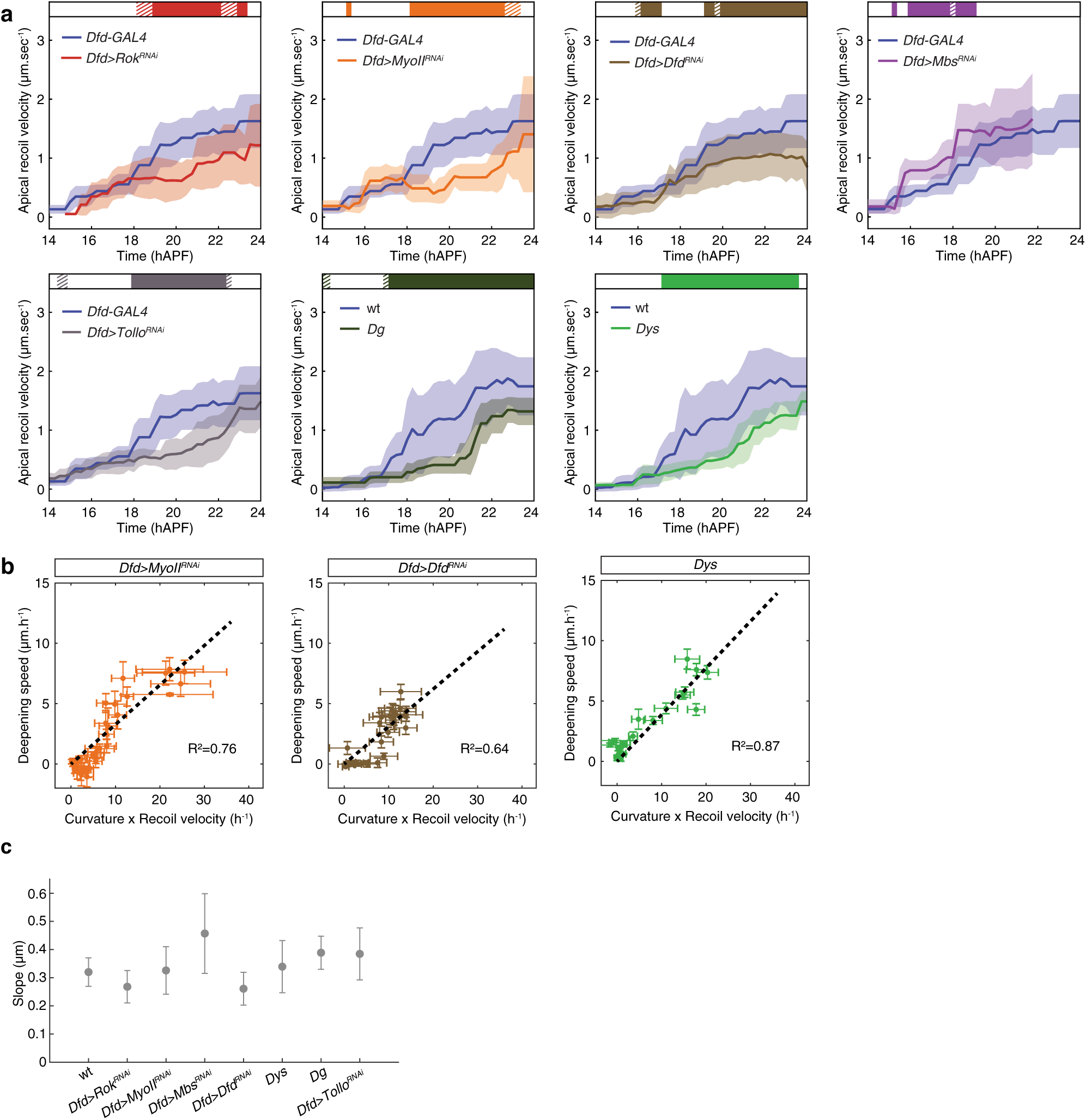
Apical recoil velocities as well as folding dynamics in wt and mutant conditions. **(a)** Graphs of the ML apical initial recoil velocity (mean ± sd, averaged with a 2h sliding window) after ablation in the neck region in control *Dfd-GAL4* (*N=*125 pupae) and wt (*N=*127 pupae) tissues, and in *Dfd*>*Rok^RNAi^* (*N=*57 pupae), *Dfd*>*MyoII^RNAi^* (*N=*48 pupae), *Dfd*>*Dfd^RNAi^* (*N=*106 pupae), *Dfd*>*Mbs^RNAi^* (*N=*61 pupae), *Dfd*>*Tollo^RNAi^* (*N=*90 pupae), *Dg* (*N=*57 pupae) and *Dys* (*N=*108 pupae) tissues. Horizontal box: *p-values* of Welch tests performed between experimental condition and either the *Dfd-GAl4* or wt control at successive time points (white *p*>0.05, striped *p*<0.05, solid *p*<0.01). **(b)** Graph of the apical fold front deepening speed and of the product of curvature and ML tension estimated by laser ablation in *Dfd>MyoII^RNAi^*, *Dfd>Dfd^RNAi^* and *Dys*. Datasets from Fig. 2e, Fig. 3a and Extended Data Fig. 8a were used. A line passing through the origin was fitted (R^2^ values are indicated). **(c)** Graph of the proportionality coefficients of deepening speed with respect to product of curvature times recoil velocity (± sd) for indicated genotypes. Datasets from Fig. 2e, Fig. 3a and Extended Data Fig. 8a were used. The slopes of the linear fit were extracted for each animal to calculate average slopes and the standard errors. Across all conditions, the slope average is 0.35µm and its standard deviation is 0.06µm.

**Extended Data Fig. 9:**
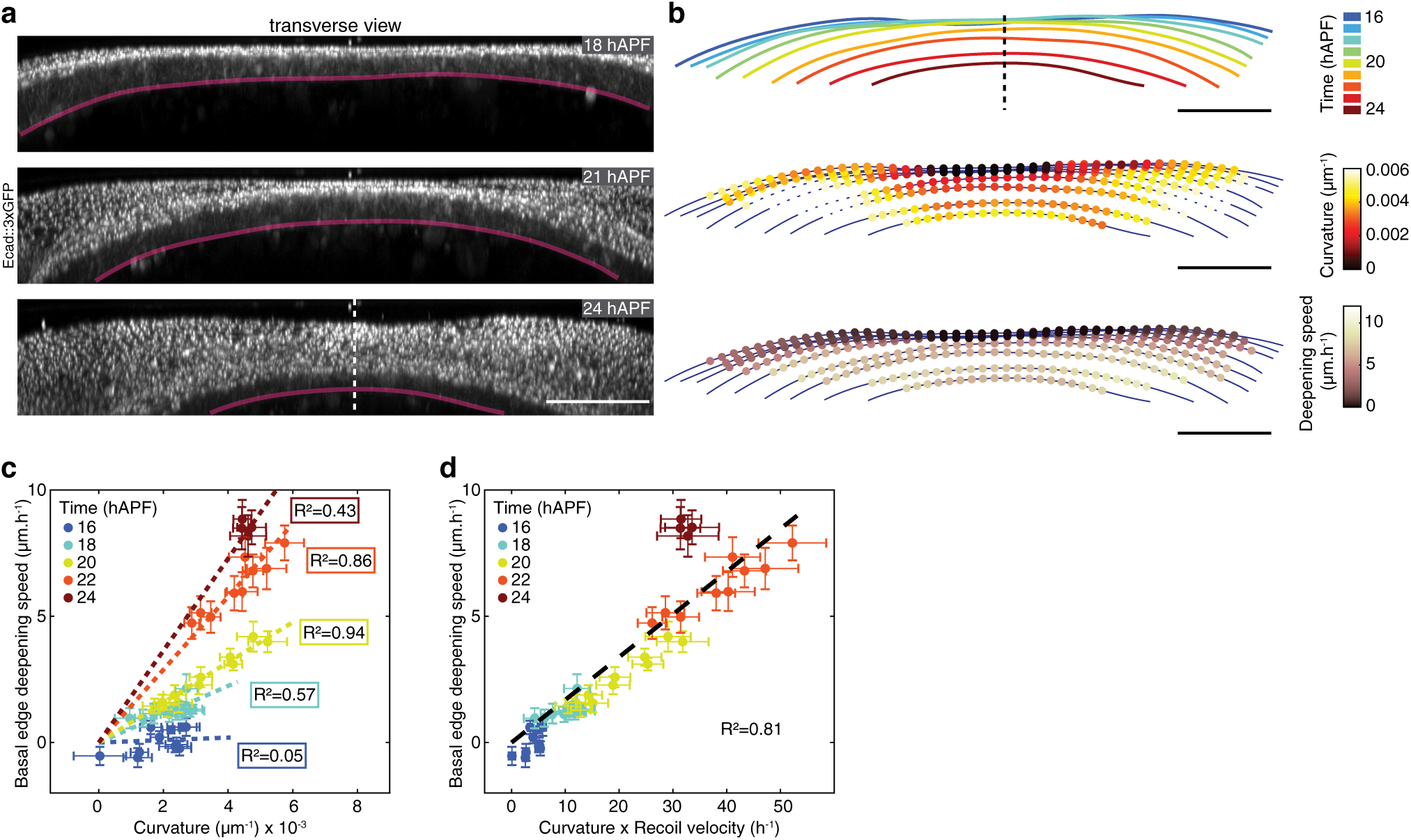
Basal folding dynamics in wt tissues. **(a)** Transverse view time-lapse images of the neck region labelled by Ecad::3xGPF at 18, 21 and 24 hAPF. Pink line, position of the basal fold front. Dashed line, midline position. **(b)** Successive positions of the basal fold front color coded according to time (top), tissue curvature (middle) and the deepening speed (bottom) for an individual pupa. Dashed line, midline position. **(c)** Graph of the basal fold front deepening speed versus curvature. Each point is the mean value (± sem) among 10 control animals for a given Position_ML_ at a given color-coded developmental time. For each developmental time, a line passing through the origin was fitted (R^2^ values are indicated). **(d)** Graph of the basal fold front deepening speed versus the product of curvature (mean ± sem) and ML basal initial recoil velocity after laser ablation in the neck region (mean ± sem) for a given Position_ML_ at a given color-coded developmental time. Datasets from Fig. 2d and Extended Data Fig. 9c were used. A line passing through the origin was fitted using average values (R^2^=0.81).

**Extended Data Fig. 10:**
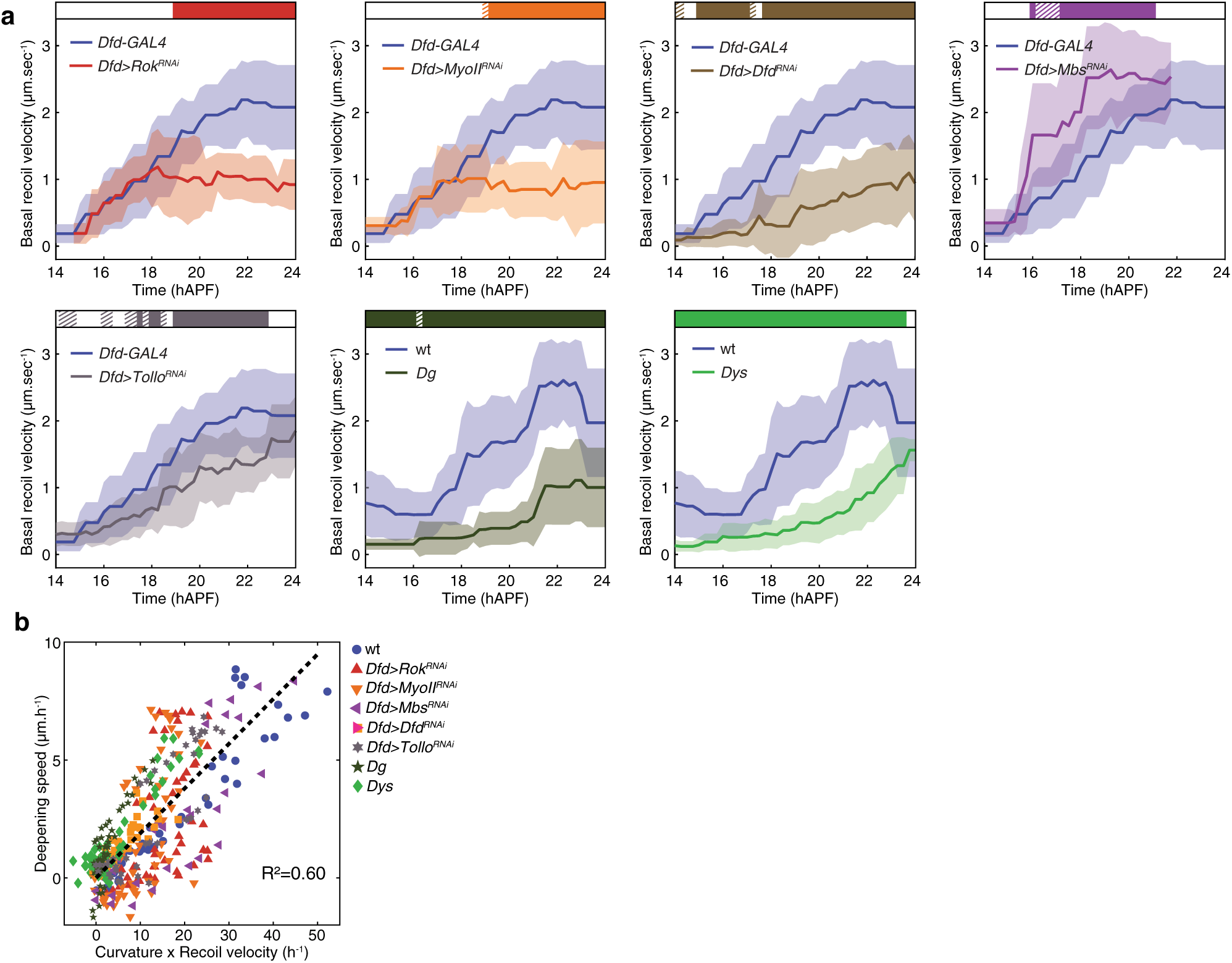
Basal recoil velocities as well as folding dynamics in wt and mutant conditions. **(a)** Graphs of the ML basal initial recoil velocity (mean ± sd, averaged with a 2h sliding window) after ablation in the neck region in control *Dfd-GAL4* (*N=*125 pupae) and wt (*N=*127 pupae) tissues, and in *Dfd*>*Rok^RNAi^* (*N=*57 pupae), *Dfd* >*MyoII^RNAi^* (*N=*48 pupae), *Dfd*>*Dfd^RNAi^* (*N=*106 pupae), *Dfd*>*Mbs^RNAi^* (*N=*61 pupae), *Dfd*>*Tollo^RNAi^* (*N=*90 pupae), *Dg* (*N=*57 pupae) and *Dys* (*N=*108 pupae) tissues. Horizontal box: *p-values* of Welch tests performed between experimental condition and either the *Dfd-GAl4* or wt control at successive time points (white *p*>0.05, striped *p*<0.05, solid *p*<0.01). **(b)** Graph of the basal fold front deepening speed (mean ± sem) and of the product of curvature and ML basal tension estimated by laser ablation (mean ± sem) for a given Position_ML_ at a given developmental time for all tested experimental conditions. Datasets used are the ones in Fig. 4f for the tracking of the basal fold front and basal recoil velocity measurements from Extended Data Fig. 9 and Extended data Fig. 10a. A line passing through the origin was fitted (R^2^=0.60).

**Extended Data Fig. 11:**
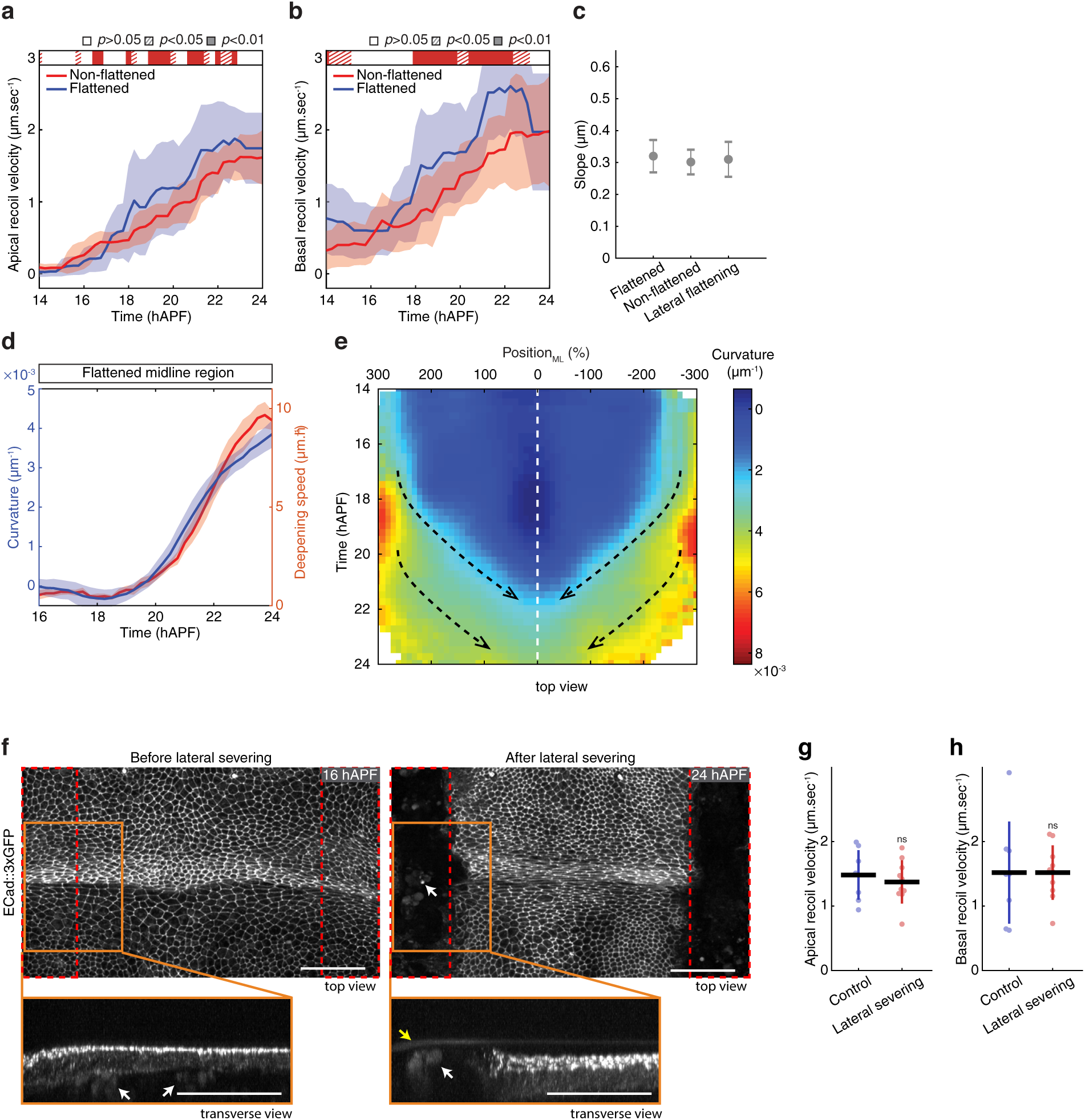
Temporal evolution of neck curvature upon flattening. **(a)** Graph of the ML apical initial recoil velocity (mean ± sd, averaged with a 2h sliding window) after ablation in the neck region in pupae with (flattened, *N=*127 pupae) and without (non-flattened, *N=*72 pupae) coverslip flattening. Horizontal box: *p-values* of Welch tests performed between flattened and non-flattened pupae at successive time points (white *p*>0.05, striped *p*<0.05, solid *p*<0.01). **(b)** Graph of the ML basal initial recoil velocity (mean ± sd, averaged with a 2h sliding window) after ablation in the neck region in pupae with (flattened, *N=*127 pupae) and without (non-flattened, *N=*71 pupae) coverslip flattening. Horizontal box: *p-values* of Welch tests performed between flattened and non-flattened pupae at successive time points (white *p*>0.05, striped *p*<0.05, solid *p*<0.01). **(c)** Graph of the proportionality coefficients of deepening speed with respect to product of curvature times recoil velocity in medially flattened, non-flattened and laterally flattened conditions (± sd). Datasets of Fig. 4d, and Fig. 5e,j were used. The slopes of the linear fit were extracted for each animal to calculate average slopes and the standard errors. Flattening affects both the curvature times tension and the velocity, but their relationship is unchanged (they remain proportional to each other with the same slope), indicating that the dissipation is unchanged, and more generally the morphogenetic mechanism remains the same. **(d)** Graph of the curvature (blue, mean ± sem) and of the deepening speed (red, mean ± sem) of the apical fold front as a function of time for the flattened midline region in 10 control animals mounted using classical mounting with coverslip (flattening). Note that the curvature gradually increases in time, which is accompanied by an increase in the deepening speed. **(e)** Color-coded evolution of the mean curvature of the apical fold front as a function of the Position_ML_ in 10 control animals mounted with coverslip flattening. Note the propagation of curvature from lateral to the medial as the apical fold front deepens and the transition from heterogenous curvature before 19 hAPF to a homogenous curvature after 22 hAPF (outlined by black dashed arrows). White dashed line, midline position. **(f)** Top view (top) and transverse view (bottom) images of Ecad::3xGFP labelling in the apical neck tissue at 16 hAPF prior to tissue severing ablation of each lateral side of the neck (red dashed boxes) and at 24 hAPF after ablation. A close up of the region outlined in orange is shown below the panel. White arrows indicate autofluorescent yolk granules. Note that these granules reside in the hemolymph underneath the epithelium prior to ablation (16 hAPF), but after ablation (24 hAPF), are near the most apical side of the tissue close to the cuticle which covers the epithelium (yellow arrow). **(g)** Graph of the ML apical initial recoil velocities (mean ± sd) upon probe ablation in the medial neck region in pupae with coverslip flattening (*N*=8) and in pupae with coverslip flattening after lateral side tissue severing ablations (*N*=10). The probe ablations to measure recoil velocities were performed at 24 ± 0.5 hAPF, ∼9h after the first lateral tissue severing ablations were performed to isolate the medial tissue region. *t*-test, ns *p*>0.05. **(h)** Graph of the ML basal initial recoil velocities (mean ± sd) upon probe ablation in the medial neck region in pupae with coverslip flattening (*N*=8) and in pupae with coverslip flattening after lateral side tissue severing ablations (*N*=10). The probe ablations to measure recoil velocities were performed at 24 ± 0.5 hAPF, ∼9h after the first lateral tissue severing ablations were performed to isolate the medial tissue region. *t*-test, ns *p*>0.05. Scale bars, 50µm.

**Supplementary Table 1.** *Drosophila* alleles and transgenes.

| *Drosophila* stock | Reference or Source |
| --- | --- |
| **Ubi-Ecad::GFP** | Ref<sup>69,83,84</sup> |
| **Ecad::3xGFP** | Ref<sup>66</sup> |
| **Ecad::3xmKate2** | Ref<sup>66</sup> |
| **utrABD::GFP** | Ref<sup>85</sup> |
| **MyoII::3xGFP** | Ref<sup>66</sup> |
| **MyoII::3xmKate2** | Ref<sup>66</sup> |
| **PH::GFP** | Ref<sup>86</sup> |
| **UAS-LifeAct::Ruby** | Bloomington Stock Center (#35545) |
| **Tollo::YFP** | Ref<sup>35</sup> |
| **Dg::GFP** | This study |
| **Dys::GFP** | Bloomington Stock Center (#59782) |
| **UAS-Diap1** | Bloomington Stock Center (#6657) |
| ***sqh*<sup>RNAds</sup>** | Bloomington Stock Center (#33892) |
| ***Rok*<sup>RNAds</sup>** | Bloomington Stock Center (#28797) |
| ***Mbs*<sup>RNAds</sup>** | Bloomington Stock Center (#41625) |
| ***Dys*<sup>RNAds</sup>** | Bloomington Stock Center (#55641) |
| ***Dfd*<sup>RNAds</sup>** | Vienna Drosophila Resource Center (v50110) |
| ***Dg*<sup>RNAds</sup>** | Vienna Drosophila Resource Center (107029) |
| ***Tollo*<sup>RNAds</sup>** | Bloomington Stock Center (#28519) |
| ***Tollo*<sup>59</sup>** | Ref<sup>87</sup> |
| ***Tollo*<sup>C5</sup>** | Ref<sup>88</sup> |
| ***Dg*<sup>086</sup>** | Ref<sup>89</sup> |
| ***Dg*<sup>043</sup>** | Ref<sup>89</sup> |
| ***Dys*<sup>E17</sup>** | Ref<sup>34</sup> |
| ***Dys*<sup>Exel6184</sup>** | Ref<sup>89</sup> |
| **hs-flp** | Bloomington Stock Center (#8862) |
| **Act>CD2>Gal4, UAS-Caax:tBFP** | Ref<sup>90</sup> |
| **Act-GAL4** | Bloomington Stock Center (#3954) |
| **Ubi-GAL80<sup>ts</sup>** | Bloomington Stock Center (#7017) |
| ***Dfd*-GAL4** | This study |

## Supplementary Video Legends

**Supplementary Video 1: Neck folding during pupal development.**

Time-lapse 3D (top) movie of the dorsal side of Ecad::3xGFP pupa acquired in the neck region (blue) and the corresponding transverse view (bottom) movie between 14 and 24h40min hAPF.

Scale bar, 50µm.

**Supplementary Video 2: Localization of Dfd, Tollo, MyoII, F-Actin and Dg during neck folding.**

**(a)** Images of Dfd (magenta) and Ecad::3xGFP (green) at 14, 18, 21 and 24 hAPF during neck fold formation. Dfd levels are constant throughout neck folding.

**(b, c)** Apical (b) and basal (c) MyoII::3xGFP (magenta) and F-Actin (green) localizations at 14, 18, 21 and 24 hAPF during neck fold formation. Apical MyoII accumulation is present in the neck cells at 14 hAPF and becomes more pronounced by 18 hAPF. From 21 hAPF onwards, MyoII gets increasingly polarized to accumulate in cell junctions (b). MyoII and F-actin are observed to form a network at the basal side of the neck at 14 hAPF. The anisotropy of this actomyosin network increases over time, becoming noticeably pronounced by 21 hAPF in the medial region of the neck (c).

**(d)** Images of Tollo::YFP (green) at 14, 18, 21 and 24 hAPF during neck fold formation. Tollo::YFP is enriched apically at neck cell junctions throughout neck invagination. From 21 hAPF onwards Tollo::YFP becomes also enriched in the lateral domains of the thorax.

**(e)** Images of Dg::GFP (green) and Dfd (magenta) at 14, 18, 21 and 24 hAPF during neck fold formation. Dg::GFP is localized in a basal network which becomes increasingly aligned in the ML direction within the neck region. At 24 hAPF, Dg::GFP accumulation in the neck is apparently lowered and a sagittal section is shown in inset for this time point to illustrate the presence of Dg::GFP at the basal side of Dfd positive neck cells. The two arrowheads indicate the positions between which the projected maximal sagittal sections were created.

Scale bar, 20µm

**Supplementary Video 3: Apical and basal laser ablation encompassing the presumptive neck region.**

Time-lapse movies after laser ablation of the apical (top) and basal (bottom) neck tissue in pupae expressing utrABD::GFP. Dashed red boxes, ablated regions.

Scale bar, 20µm.

**Supplementary Video 4: Dynamics of neck folding as function of neck region curvature.**

Transverse view time-lapse movies of the neck region labelled by Ecad::3xGFP in pupae imaged with medial coverslip flattening (top), without coverslip flattening (middle), and with lateral coverslip flattening (bottom). The positions of the apical fold front are color-coded at 16, 18, 20, 22 and 24 hAPF on the last images.

Scale bar, 50µm.

**Supplementary Video 5: Dynamics of neck folding after lateral tissue severing ablations in the presence or the absence of medial curvature.**

Transverse view time-lapse movies of the neck region labelled by Ecad::3xGFP in a control pupa imaged with coverslip flattening (top) and in pupae with lateral tissue severing ablations and imaged without (middle) and with coverslip flattening (bottom). The positions of the apical fold front are color-coded at 16, 18, 20, 22 and 24 hAPF on the last images. Red dashed boxes, ablated regions.

Scale bar, 50µm.

## Notes

### Competing Interest Statement

The authors have declared no competing interest.

### Summary of Updates

Minor modifications to the text and the reference list

